# rsCaMPARI: an erasable marker of neuronal activity

**DOI:** 10.1101/798298

**Authors:** Fern Sha, Ahmed S. Abdelfattah, Ronak Patel, Eric R. Schreiter

## Abstract

Identifying and comparing active neuron ensembles underlying complex behaviors is a key challenge in neuroscience. Recent tools such as CaMPARI have enabled the optical marking and selection of active neuron populations. However, CaMPARI photoconversion is permanent and irreversible, limiting its utility when multiple activity snapshots must be compared within the same sample. We sought to overcome these limitations by developing an erasable neuronal activity marker based on a reversibly switchable fluorescent protein. Here we introduce rsCaMPARI, a reversibly switchable calcium marker that enables spatiotemporally precise marking, erasing, and remarking of active neuron populations under brief, user-defined time windows of light exposure. rsCaMPARI photoswitching kinetics are modulated by calcium concentration when illuminating with blue light, and the fluorescence can be reset with violet light. We demonstrate the utility of rsCaMPARI for marking and remarking active neuron populations in freely-swimming zebrafish.

## Introduction

Important behaviors such as response to stimuli and memory retrieval are governed by patterns of coordinated neuron activity in the brain. Identifying which neuron ensembles are involved and how they compare across different behavior states is critical for our understanding of brain circuitry and function. Traditional methods for identifying active neuron ensembles have relied on exploiting the cellular machinery that control expression of immediate early genes (IEGs) such as *Egr1/Zif268*, *c-Fos*, or *Arc*^*1–5*^. However, the temporal resolution of these methods is poor, typically on the scale of tens of minutes to hours, and they do not correlate well with electrical activity^6,7^. Calcium transients, in contrast, follow electrical activity more quantitatively^8,9^. Genetically encoded calcium indicators such as GCaMPs^10–12^ have enabled the visualization of calcium transients with fast kinetics, but their requirement for constant monitoring limit the field of view size, and recording neural activity in freely moving specimens is challenging. An alternative approach is to mark the active neuron ensemble during a behavior of interest for post-hoc analysis. These methods often couple calcium activity with light activation in order to drive a fluorescence change or to activate transcription factors^13,14^.

Recently we introduced CaMPARI^15^ (calcium-modulated photoactivatable ratiometric integrator) and its improved variant CaMPARI2^16^, a genetically encoded fluorescent sensor that enables spatiotemporally precise labelling of active neuron ensembles in large tissue volumes under widefield illumination. CaMPARI is a photoconvertible FP where efficient green-to-red photoconversion only occurs at elevated intracellular calcium concentration and is gated by user-defined violet light illumination. CaMPARI therefore allows integration and post-hoc analysis of the evoked calcium activity as an accumulated red signal during a specific epoch of interest, typically on the order of seconds. CaMPARI photoconversion is irreversible, which is useful when a permanent signal is desired; however, this irreversibility also hinders reuse of the tool, especially when multiple snapshots of activity are desired from the same sample preparation. This can be important when comparing active neuron ensembles in non-stereotyped organisms where there is variability in neural circuitry across different individuals, or where there is variability from trial to trial in the same individual. Reuse of CaMPARI requires waiting for protein turnover of the sensor, which may take days or weeks. An erasable probe would instead allow multiple snapshots of neuronal activity to be captured quickly within the same sample preparation.

Reversibly switchable fluorescent proteins can be toggled between a fluorescent state (on) and a non-fluorescent state (off) depending on the wavelength of light used for illumination^17^. For example, Dronpa^18^, rsEGFP^19^, and mGeos^20^ are bright green fluorescent proteins that can be off-switched with blue light illumination. Subsequent illumination with violet light reverts them back to the on-state. The structural mechanism underlying this reversible switching behavior is a cis/trans isomerization of the chromophore, which is coupled to the chromophore’s protonation state^21,22^.

We hypothesized that reversible photoswitching could provide a mechanism for marking, erasing, and remarking calcium activity using light. Specifically, we aimed to engineer an erasable calcium marker such that the photoswitching kinetics are Ca^2+^-dependent and that the final magnitude of fluorescence change provides a stable but erasable readout for the relative amount of calcium activity during a user-defined time window of light illumination; reverse photoswitching could then quickly erase the signal and reset the tool for immediate reuse. In addition, we sought to use blue light as the gate to record calcium activity, which avoids the more phototoxic near-UV light that was required for CaMPARI^23,24^.

Here we report development of a new type of erasable calcium marker called rsCaMPARI (reversibly switchable CaMPARI)(Fig. 1a). Using directed evolution methods, we engineered the off-switching kinetics of rsCaMPARI under blue light illumination to be Ca^2+^-dependent, which allows reversible calcium activity marking with cellular resolution. The fluorescence can be easily recovered with violet light for immediate reuse and enables erasable and repeatable calcium activity marking over practical timescales of a few seconds. We demonstrate the utility of rsCaMPARI for marking, erasing, and re-marking active neuron ensembles in primary neuron cultures and freely-swimming zebrafish.

**Fig 1.**
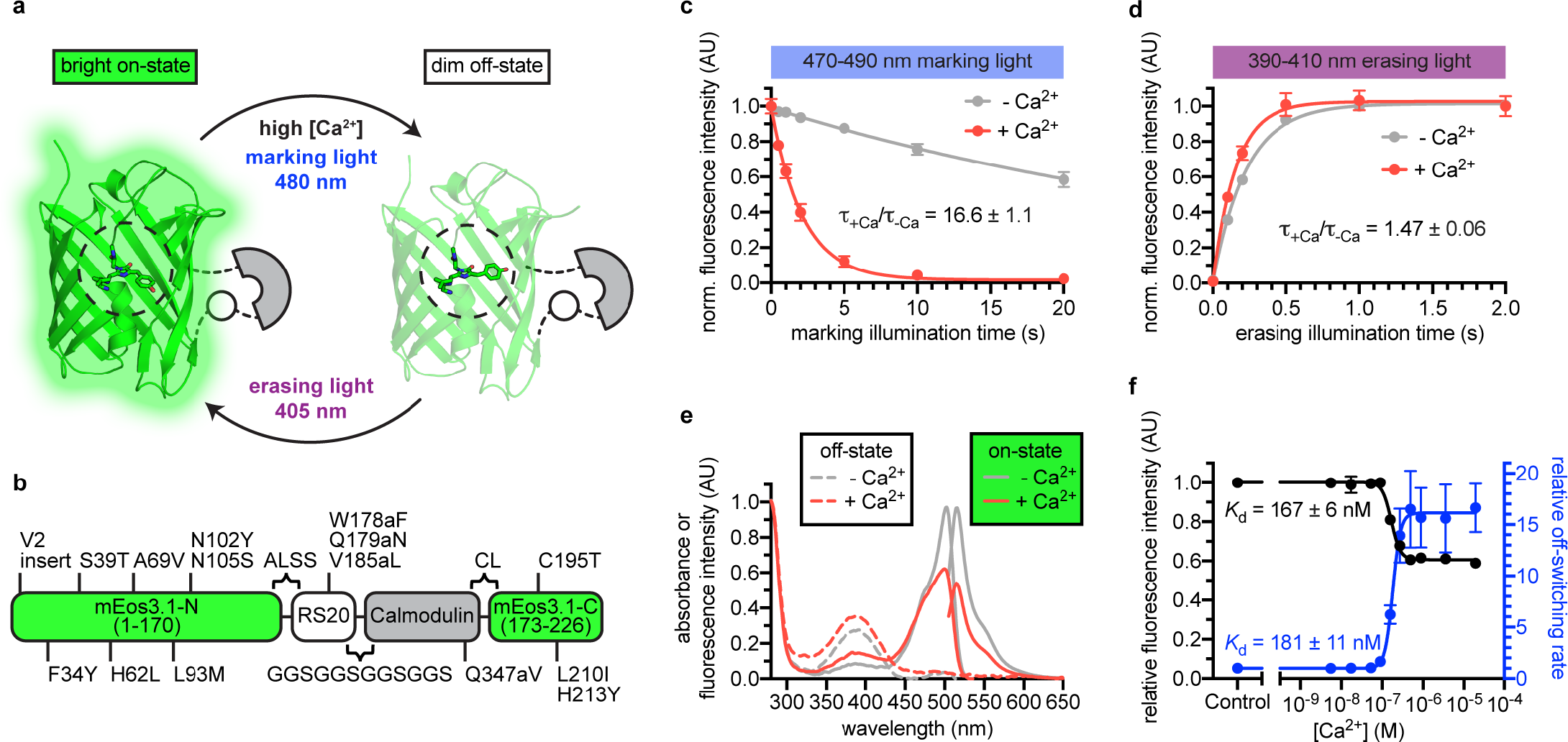
Engineering and *in vitro* characterization of rsCaMPARI. **a**, Schematic of rsCaMPARI function. **b**, Primary structure of rsCaMPARI relative to mEos3.1. **c**, Off-switching time-course of rsCaMPARI under marking light illumination (200 mW/cm^2^) in the presence or absence of calcium. Lines are single-exponential fits to data. Error bars are standard deviation, n=4 replicate measurements. **d**, On-switching time-course of rsCaMPARI under erasing light illumination (200 mW/cm^2^) in the presence or absence of calcium. Lines are single-exponential fits to data. Error bars are standard deviation, n=4 replicate measurements. **e**, Spectral properties of rsCaMPARI. Absorbance or fluorescence emission spectra of rsCaMPARI in the fluorescent on-state in the presence or absence of calcium shown with solid traces. Absorbance spectra of rsCaMPARI in the non-fluorescent off-state in the presence or absence of calcium shown with dashed traces. **f**, Relative fluorescence intensity and relative off-switching rate of rsCaMPARI as a function of free [Ca^2+^]. Lines are sigmoidal fits to data. Error bars are standard deviation, n=4 replicate measurements.

## Results

### *Engineering and* in vitro *characterization of rsCaMPARI*

To engineer an erasable calcium marker, we took inspiration from CaMPARI and CaMPARI2 since they were previously shown to function well in several *in vivo* preparations^15,16^. We introduced CaMPARI2 mutations onto a mEos3.1^25^ scaffold and converted the protein from a green-to-red photoconvertible FP to a reversibly switchable FP that cycles between bright green and dim states via an H62L substitution within the chromophore, as was previously demonstrated by the production of the mGeos FPs from EosFP^20^. After photoswitching to the off-state, the H62L mutant exhibited slow spontaneous recovery to the on-state with a half-time of several hours^20^. We reasoned that slow spontaneous recovery would be beneficial for stable and precise readout of the final fluorescence signal following photoswitching.

We next engineered libraries of insertions of a calcium-binding domain (calmodulin) and calmodulin-binding peptide into the H62L-substituted FP (Supplementary Fig. 1). Guided by crystal structures of CaMPARI^15^ and the cis/trans isomers of IrisFP^26^, we targeted the insertion of calcium-binding domains to the middle of β-strands 8 and 9 to efficiently propagate Ca^2+^-induced conformational changes to amino acids around Y63 of the chromophore, which undergoes cis/trans isomerization during reversible photoswitching. The libraries included both possible orientations of the calcium-binding domains, and had diversity in the length and composition of linkers connecting the calcium-binding and FP domains. Screening in *E. coli* lysate (Supplementary Fig. 2), we aimed to optimize four parameters: (1) difference in green fluorescence following 490nm light illumination +/− Ca^2+^, (2) recovery of fluorescence intensity following 400 nm light illumination, (3) minimum fluorescence change due to calcium binding in the absence of light illumination (“indicator behavior”), and (4) green brightness. 96 clones were selected and sequenced, resulting in 19 unique clones that exhibited a variety of photoswitching kinetics (Supplementary Table 1).

One variant with an insertion of calcium-binding domains in β-strand 9 was selected for further characterization due to its >10-fold photoswitching rate contrast, relatively small indicator behavior, and high brightness. We named this variant rsCaMPARI (reversibly switchable CaMPARI) (Fig. 1, Supplementary Fig. 3, and Supplementary Fig. 4). Additional site-saturation mutagenesis to improve rsCaMPARI contrast did not identify improved variants without compromising other desired traits. In particular, we observed that any improved fluorescence contrast appeared to be tightly coupled to increased indicator behavior (Supplementary Fig. 5).

rsCaMPARI is a bright green and calcium-dependent photoswitchable FP (Fig. 1, Table 1, and Supplementary Fig. 6). During illumination with blue “marking light” (470-490 nm, 200 mW/cm^2^, Supplementary Fig. 7e), rsCaMPARI exhibited ~17-fold faster off-switching kinetics in high calcium compared to low calcium (Fig. 1c) and the off-switching rate was linear with respect to the light power (Supplementary Fig. 8). We observed a strong dependence of the extent of photoswitching and the rate contrast on the wavelength of light applied (Supplementary Fig. 7). When exposed to violet “erasing light” (390-410 nm, 200 mW/cm^2^), the recovery of rsCaMPARI fluorescence back to the bright on-state was very efficient and had similar on-switching kinetics in either high or low calcium conditions (Fig. 1d). Measurement of green fluorescence intensity as a function of free calcium concentration showed that the calcium free state was ~2-fold brighter than the calcium-bound state (Fig. 1e,f), and indicated an apparent dissociation constant of 167 ± 6 nM (Fig. 1f). Measurement of the off-switching rate vs. free calcium concentration gave a similar apparent dissociation constant of 181 ± 11 nM (Fig. 1f). The calcium affinity of rsCaMPARI is therefore similar to previously described CaMPARIs^15,16^ and GCaMPs^10–12^, suggesting that the dynamic range of rsCaMPARI calcium sensitivity falls within the physiological range of neuronal calcium transients.

**Table 1.** Photophysical properties of rsCaMPARI.

| | Ca<sup>2+</sup> | $\lambda_{\text{abs}}$ (nm) | $\lambda_{\text{ex}}$ (nm) | $\epsilon$ (M<sup>-1</sup>cm<sup>-1</sup>) | $\Phi_F$ | Brightness<sup>a</sup> | ↓ rate<sup>b</sup> (s<sup>-1</sup>) | ↑ rate<sup>c</sup> (s<sup>-1</sup>) | $K_d$ (nM) | Hill coefficient |
| --- | --- | --- | --- | --- | --- | --- | --- | --- | --- | --- |
| rsCaMPARI (on) | - | 502 | 515 | 60304 | 0.54 | 32.6 | 0.028 ± 0.002 | - | 167 ± 6 | 3.96 |
|  | + | 500 | 515 | 38234 | 0.46 | 17.6 | 0.467 ± 0.066 | - |  |  |
| rsCaMPARI (off) | - | 392 | - | - | - | - | - | 4.37 ± 0.24 | - | - |
|  | + | 389 | - | - | - | - | - | 6.40 ± 0.28 |  |  |

<sup>a</sup>Brightness = $\epsilon \times \Phi_F / 1000$
<sup>b</sup>measured during irradiation with 200 mW/cm<sup>2</sup> of 484 nm light
<sup>c</sup>measured during irradiation with 200 mW/cm<sup>2</sup> of 405 nm light

### Characterization of rsCaMPARI in dissociated neurons

We expressed rsCaMPARI in dissociated primary rat hippocampal neurons fused to a mRuby3^27^ fluorescent tag to normalize for expression level. We used field electrode stimulation to induce action potential firing at 80 Hz during continuous illumination with marking light (200 mW/cm^2^) on an epifluorescence microscope (Fig. 2a-e and Supplemental Fig. 9). Neurons illuminated with 10 s of marking light accompanied by field stimulation exhibited strong dimming of the green fluorescence signal (Fig. 2a, cycle 1), which could later be recovered by a 3-second pulse of erasing light. When the same neuron was subsequently illuminated with marking light but without field stimulation, only modest dimming of the green fluorescence was observed (Fig. 2a, cycle 2). The same neuron could be repeatedly marked and erased over several cycles (Fig. 2a, cycles 3-6). We observed 3-fold contrast in rsCaMPARI fluorescence between stimulated and non-stimulated neuron cultures after exposure to marking light (Fig. 2b). Monitoring green fluorescence of the neurons over time showed a shift to more rapid off-switching that correlated with the onset of field stimulation (Fig. 2c). In response to three trains of 160 action potentials during marking light, we observed the largest fluorescence change during the first train (~40-60% green fluorescence decrease compared to ~10% in non-stimulated neurons). Smaller fluorescence changes were observed during the second and third field pulse trains as the sensor approached near complete off-switching.

**Fig 2.**
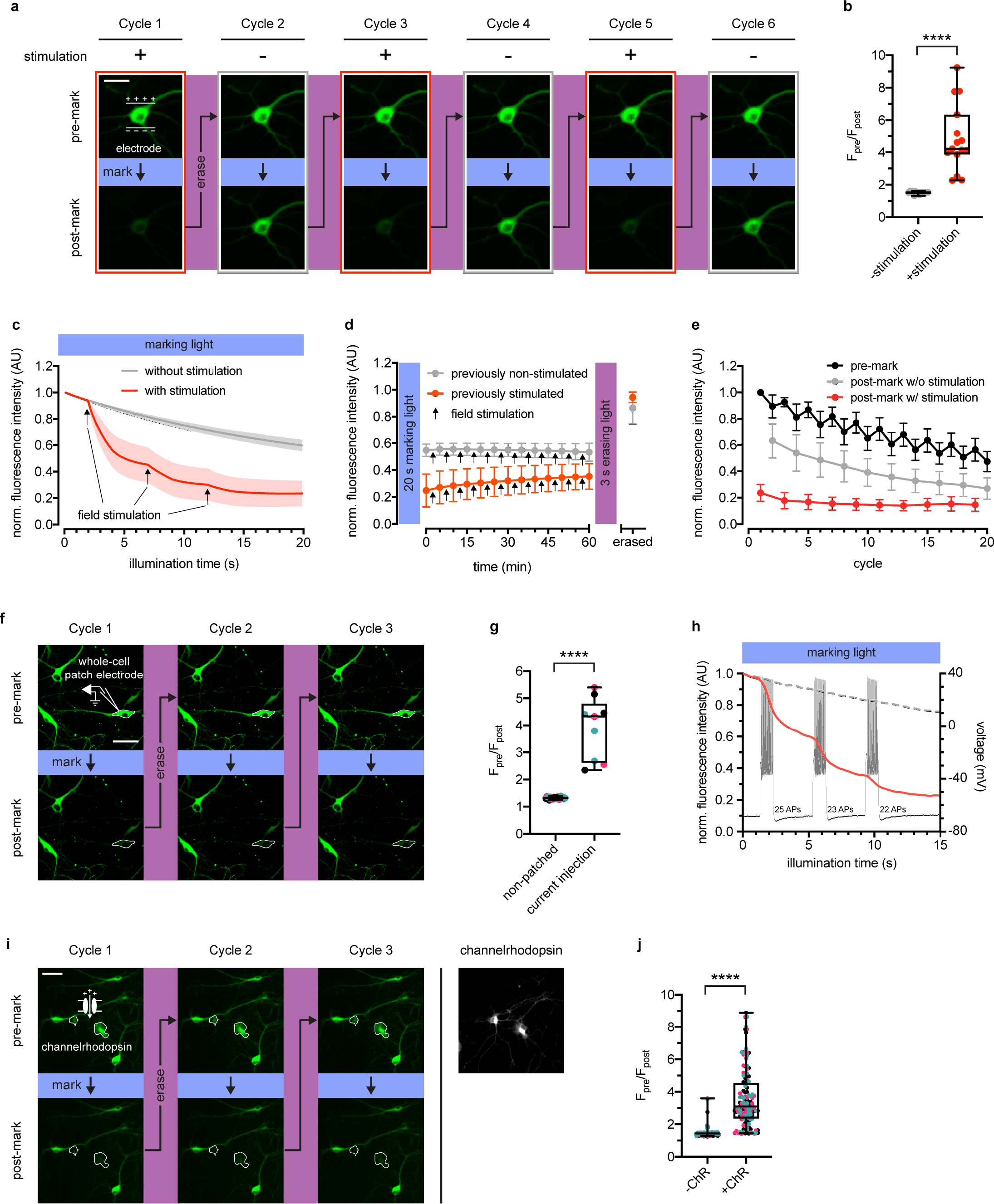
rsCaMPARI reversibly and selectively marks activity in primary neurons. **a**, Fluorescence images of a representative primary rat hippocampal neuron undergoing multiple cycles of exposure to a 10 s window of marking light (224 mW/cm^2^) +/− field stimulation (2x 160 stimulations at 80 Hz). Each cycle was reset with a 3 s pulse of erasing light (224 mW/cm^2^). Scale bar is 30 μm. **b**, Quantification of pre-marked to post-marked green fluorescence ratios of individual neurons (n=15 neurons from two independent wells) +/− field stimulation (2x 160 stims at 80 Hz). Boxplot whiskers extend from min to max values and box extends from 25^th^ to 75^th^ percentile. Middle line in box is the median. ****P < 0.0001, two-tailed Student’s *t*-test. **c**, Fluorescence time-course traces of neurons undergoing one cycle of illumination +/− stimulation (3x 160 stims at 80 Hz). Arrows on time-course trace denote start of each stimulation. Error bars are standard deviation, n=15 neurons from two independent wells. **d**, Time-course of rsCaMPARI spontaneous recovery in the dark at 37°C following marking light illumination of previously non-stimulated or previously stimulated neurons. Arrows denote a bout of field stimulation between each imaging timepoint (160 stims at 80 Hz). After one hour, the neurons were reset with a 3 s pulse of erasing light. Error bars are standard deviation, n=66 previously non-stimulated neurons and n=59 previously stimulated neurons from two independent wells. **e**, Photofatigue of rsCaMPARI over successive cycles of marking light illumination with or without field stimulation. Each cycle is followed by erasing light to reset the sensor. Error bars are standard deviation, n=7 neurons from three independent wells. **f**, Fluorescence images of rsCaMPARI before and after 15 s of marking light illumination (150 mW/cm^2^). A single cell, denoted by pipette drawing, was patched and stimulated to fire action potentials by injecting current during marking light illumination. Scale bar is 50 μm. **g**, Quantification of pre-marked to post-marked green fluorescence ratios of individual neurons across three marking cycles for patched (n=3 neurons from three independent wells) and non-patched cells (n=8 neurons from three independent wells). Cyan, red, and black data points are measurements from the first, second, and third cycles, respectively. ****P < 0.0001, two-tailed Student’s *t*-test. **h**, Single-trial recording of action potentials from current injection during the first cycle in patched neuron shown in (f) using fluorescence imaging (red trace) or electrophysiology to measure membrane potential (black trace). Average fluorescence traces of non-patched neurons are shown as dashed grey trace. **i**, Fluorescence images of rsCaMPARI before and after 10 s of marking light illumination (285 mW/cm^2^)(left panels). Two neurons denoted by a white outline are positive for channelrhodopsin (ChR) expression. Right-most panel is a fluorescence image of channelrhodopsin expression. Scale bar is 100 μm. **j**, Quantification of pre-marked to post-marked green fluorescence ratios of individual neurons across three marking cycles for +ChR (n=42 neurons from 17 independent wells) and -ChR (n=79 neurons from 17 independent wells) cells. Cyan, red, and black data points are measurements from the first, second, and third cycles, respectively. ****P < 0.0001, Wilcoxon rank-sum test.

In order for rsCaMPARI activity marking to be compatible with post-hoc analysis, it is important that the post-marking fluorescence is stable. We tracked the fluorescence of neurons that had previously been illuminated with marking light and found that rsCaMPARI green fluorescence is stable in the dark for more than one hour (Fig. 2d, Supplementary Fig. 9c, and Supplementary Fig. 10), even during ongoing calcium activity. This provides sufficient time for readout of the recorded signal, for example with high-resolution microscopy techniques over large volumes of tissue, and demonstrates that rsCaMPARI is sufficiently stable for post-hoc analysis.

Next, we characterized photofatigue of rsCaMPARI in neurons from multiple rounds of marking and erasing to understand how many times activity patterns could be marked. Fluorescence was monitored before and after marking light illumination, then recovered with erasing light over successive cycles (Fig. 2e). The neurons were stimulated during odd-numbered cycles, but not during even-numbered cycles. We observed that rsCaMPARI lost ~50% of its initial fluorescence after 20 cycles. The fluorescence contrast between “stimulated” and “not stimulated” was reduced to ~50% of the initial difference after 10 cycles, but still allowed discrimination between marked and unmarked cells. rsCaMPARI may therefore be erased and reused for at least 10 cycles of marking activity.

To demonstrate that rsCaMPARI can mark an active subset of neurons within a population, we first performed whole-cell patch clamp electrophysiology on a single rsCaMPARI-expressing neuron, delivering current injections to produce controlled action potential firing during marking light (Fig. 2f-h and Supplementary Fig. 11). Other neurons in the same field of view were not active due to application of drugs to block synaptic release. The patched neuron became much dimmer during the marking light illumination (Fig. 2f, bottom panels), providing 4-fold contrast when compared to surrounding neurons that were not patched (Fig. 2g), and more rapid off-switching correlated well with action potential firing (Fig. 2h). The green rsCaMPARI fluorescence could be quickly recovered by subsequent illumination with erasing light and selective marking of the patched neuron could be repeated several times (Fig. 2f, cycles 2 and 3). We also drove activity in a subset of neurons using a channelrhodopsin. We sparsely co-expressed ChrimsonR^28^ channelrhodopsin in neurons expressing rsCaMPARI and acquired images before and after illumination with marking light and pulsed 560 nm light to fully drive the channelrhodopsin (Fig 2i-j and Supplementary Fig. 12). A subset of neurons became much dimmer (Fig. 2i, bottom panels), primarily corresponding to neurons that were positive for channelrhodopsin (Fig. 2i, right panel; Supplementary Fig. 12a, bottom panels) and showed 3-fold contrast (Fig. 2j). The same subset of neurons could be repeatedly erased and marked again (Fig. 2i, cycles 2 and 3).

### Marking activity patterns in freely-swimming zebrafish

To demonstrate the utility of rsCaMPARI for marking active neurons *in vivo*, we generated stable transgenic zebrafish (*Danio rerio*) expressing rsCaMPARI from a neuron-specific promoter (Fig. 3a). We developed an imaging protocol for rsCaMPARI in the zebrafish brain using light sheet fluorescence microscopy for rapid acquisition and to minimize out-of-plane excitation light (Fig. 3b). Zebrafish larvae (4-5 days post fertilization) experiencing a variety of stimuli were exposed to 10 s of marking light (400 mW/cm^2^) and light sheet z-stacks were acquired in the following order: (1) acquire the post-marking signal as a “marked” stack, (2) reset to the bright on-state with erasing light, and (3) acquire the post-erased signal as a “reference” stack. Comparison between the marked and reference stacks thus revealed the relative magnitude of rsCaMPARI photoswitching, which we quantitated and normalized as a ΔF/F metric. When the fish were anesthetized with the sodium channel blocker tricaine methanosulfonate (MS-222) to block brain activity during exposure to marking light, there was low and uniform labeling across the brain, consistent with a small amount of background photoswitching in the absence of calcium (Fig. 3c and Supplementary Fig. 13). However, in freely-swimming zebrafish, we observed neuron-specific labeling patterns, particularly in the forebrain, habenula, and hindbrain, which were consistent with labeling patterns observed previously using CaMPARI^15^. Additional stimulation with cold or warm water, or the proconvulsant potassium channel blocker 4-aminopyridine (4-AP) also produced distinct and reproducible patterns of labeling.

**Fig 3.**
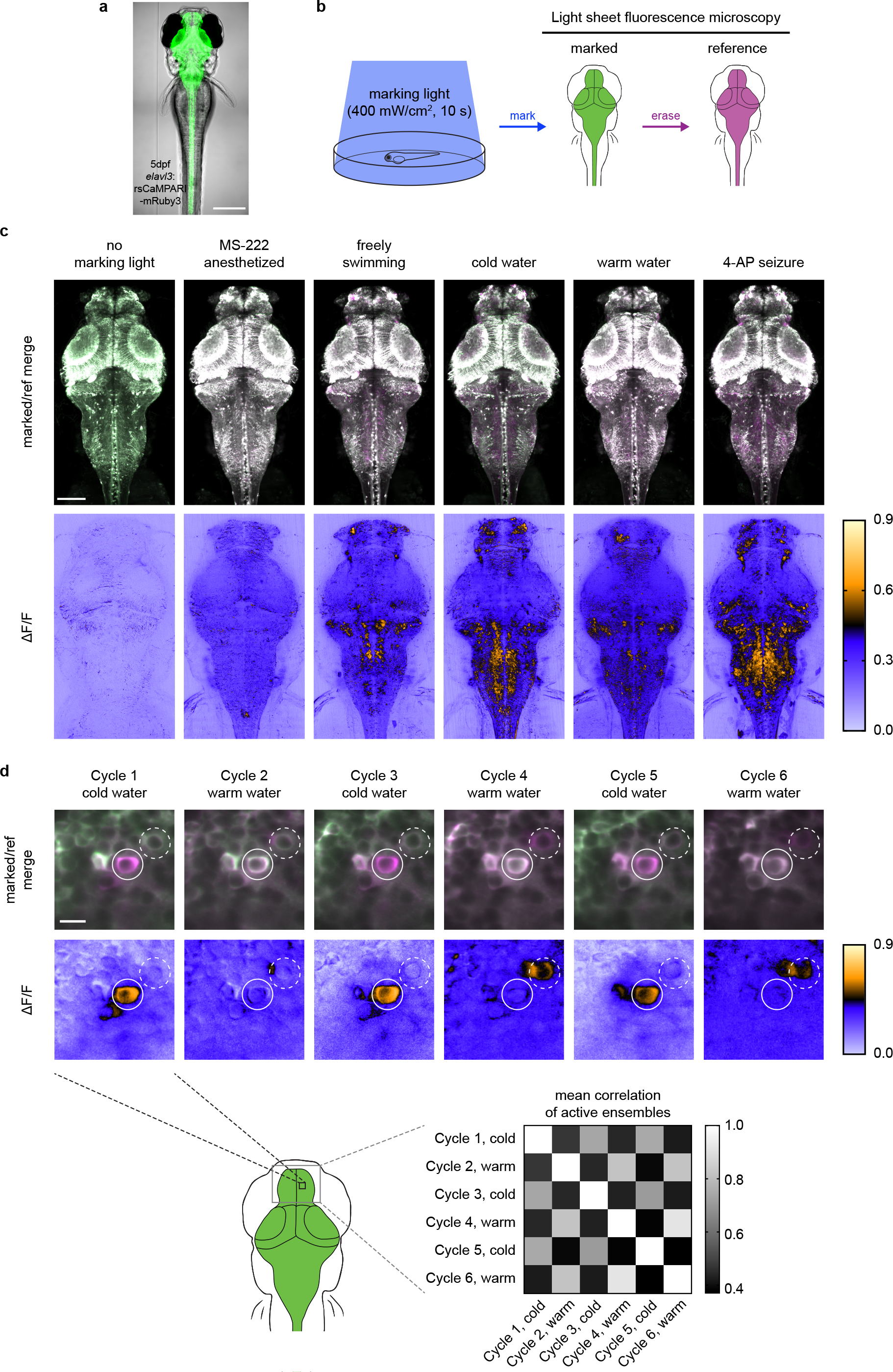
rsCaMPARI reversibly marks active neuron ensembles in the freely moving larval zebrafish. **a**, rsCaMPARI expression in the Tg[*elavl3:*rsCaMPARI-mRuby3]^jf93^ zebrafish. Scale bar is 300 μm. **b**, Cartoon schematic of experimental setup and image acquisition. **c**, Maximum intensity Z projections of the entire brain from zebrafish larvae (4 to 5 dpf) after exposure to different stimuli: no marking light, anesthetized with tricaine methanesulfonate (MS-222), freely swimming in system water, cold water (4°C), warm water (45°C), or 4-aminopyridine (4-AP). Top panels are merged reference and erased images, pseudo-colored green and magenta, respectively. Bottom panels are ΔF/F images. Scale bar is 100 μm. **d**, Multiple cycles of rsCaMPARI marking in the same zebrafish (5 dpf) exposed to either cold or warm water. Top panels are individual Z slices from the pallium of the same fish brain illustrating one neuron (solid white circle) that was only labeled during cold stimulus and another neuron (dashed white circle) that was variably labeled during warm stimulus. Scale bar is 10 μm. Bottom-right panel is a mean correlation matrix comparing active ensembles in the upper pallium and habenula across multiple marking cycles in the same fish.

Finally, to demonstrate the ability of rsCaMPARI to reproducibly mark multiple distinct activity patterns *in vivo*, we performed multiple marking and erasing cycles in the same fish during alternating exposures to cold and warm water (Fig. 3d and Supplementary Fig. 14). We observed stimulus-specific labeling across multiple marking cycles for neurons located in the zebrafish pallium. For example, over six marking cycles (alternating cold and warm water), specific neurons were strongly and repeatedly labeled during cold water exposures, but not during warm water stimulus (Fig. 3d, upper panels). We also observed some variability across trials when using the same stimulus, presumably due to variability in the ongoing brain activity leading up to the stimulus and the variability in the perception to the stimulus experienced by freely swimming fish. Overall, the patterns of labelling across the pallium in response to either cold or warm water were specific to the stimulus: activity patterns in response to cold water showed higher mean correlation with other cold response patterns, and activity patterns in response to warm water showed higher mean correlation with other warm response patterns (Fig. 3d, bottom-right panel). Taken together, these results demonstrate that rsCaMPARI is able to consistently capture active neuron ensembles from specific stimuli over multiple cycles *in vivo*.

## Discussion

We recently introduced CaMPARI^15,16^, a fluorescent protein whose permanent green-to-red photoconversion is calcium-dependent, to enable marking of active neurons with finer time resolution than activity-dependent gene expression. CaMPARI has been successfully used for marking active neuronal circuits, but is limited by the irreversible nature of its color change, making it essentially a one-shot reporter. Assignment of ensembles of neurons to specific functional circuits is greatly facilitated by comparisons between stimulus and control periods or between repeated trials, which can’t be achieved with a single snapshot. Our experiments in zebrafish demonstrate how rsCaMPARI now enables marking of multiple active ensembles of neurons by allowing the probe to be reset by erasing light between trials. Thus, the erasable nature of rsCaMPARI now allows comparisons to be made between activity patterns across multiple trials from a variety of different stimuli, all within the same sample preparation.

We achieved erasable marking of neuronal activity by engineering reversibly switchable fluorescent proteins, a class of fluorescent probes that has been utilized for superresolution microscopy approaches but which has not yet been exploited to create functional activity reporters. Another possible future application of reversibly switchable activity reporters is their use with superresolution microscopy techniques like nonlinear structured illumination to visualize physiological signals within intracellular microdomains.

In summary, rsCaMPARI retains many of the advantages of CaMPARI such as bright fluorescence, high contrast, light gating, short integration windows (seconds), and stable integrated signal, on top of being reusable. Additionally, rsCaMPARI can be used in a single (green) fluorescent color channel by comparing the post marking light image to the post erasing light image as we show in zebrafish, allowing the simultaneous use of complementary reporters or effectors in other color channels. We believe rsCaMPARI represents an important addition to the toolkit for marking active neuronal populations and we expect it will enable functional circuit marking and mapping with higher accuracy than previously possible.

## Acknowledgements

We thank members of the Janelia Experimental Technology (jET), Media Prep, Molecular Biology, Viral Tools, Vivarium, and Light Microscopy cores. Specifically, we wish to thank Damien Alcor, Michael DeSantis, Benjamin Foster, Vasily Goncharov, Cameron Loper, Igor Negrashov, Kimberly Ritola, Jared Rouchard, Steven Sawtelle, Jordan Towne, Deepika Walpita, and Xiaorong Zhang. We thank Kaspar Podgorski for comments on the project and manuscript.

## Author contributions

F.S. and E.R.S. conceived the project. F.S. engineered rsCaMPARI. F.S., A.S.A., and R.P. performed *in vitro* experiments. F.S. performed experiments in larval zebrafish. F.S. and E.R.S. analyzed data. F.S. and E.R.S. wrote the manuscript, with contributions from all other authors.

## Competing Interests

F.S. and E.R.S are listed as inventors on a patent application describing reversibly switchable neuronal activity markers.

## Materials and Methods

### Reagent availability

DNA constructs for pRSET_His-rsCaMPARI-mRuby3, pAAV-hsyn_NES-His-rsCaMPARI-mRuby3, pAAV-hsyn_NLS-His-rsCaMPARI-mRuby3, and pTol2-elavl3_NES-rsCaMPARI-mRuby3 are available via Addgene (http://www.addgene.org #120804, #120805, #122092, and #122129, respectively). Tg[*elavl3*:rsCaMPARI-mRuby3]^jf93^ transgenic zebrafish are deposited to the Zebrafish International Resource Center (https://zebrafish.org).

### Directed evolution of rsCaMPARI

CaMPARI2 in a pRSET plasmid (Life Technologies) was circularly permutated back to the original N- and C-termini of EosFP. The CaMPARI2 calmodulin and RS20 peptide were deleted to repair the original β-strand 7 of mEos3.1. H62 of the chromophore was mutated to L with a mutagenic primer using the QuikChange method (Agilent) to produce a reversibly switchable mEos3.1 (rs-mEos3.1) construct variant. This construct contained the following mutations (derived from CaMPARI) relative to mEos3.1: V2 insert, F34Y, S39T, H62L, A69V, L93M, N102Y, N105S, C195T, L210I, and H213Y. A red fluorescent protein, mCherry^29^, was fused in frame at the C-terminus to normalize for expression and photoswitching. The plasmid was linearized by PCR such that calcium-binding domains could be inserted into β-strand 8 or 9. Calmodulin and RS20 were fused with a flexible (GGS)_4_ amino acid linker in either the CaM-RS20 or RS20-CaM orientation for insertion as a single fragment using the Gibson assembly method^30^. The fragment was amplified with primers that contained (1) a variable region with 2 NNS codons plus 0 to 4 codons originating from EosFP in order to introduce linker composition and length diversity, and (2) a 5’ 30 bp region for annealing to the linearized plasmid. Each β-strand library had a theoretical diversity size of 8 × 10^6^. The Gibson assembly mixtures were diluted 3-fold in deionized water and transformed into T7 express *E. coli* (New England Biolabs) by electroporation. Transformed bacteria were plated on LB + ampicillin to isolate single colonies and a small number of clones were sequenced to confirm the desired library diversity.

Fluorescent colonies were manually picked and transferred to liquid growth media in 96-deep well plates and grown and harvested as previously described^15^. Two wells containing the rs-mEos3.1 construct without calcium-binding domain insertions and two wells containing a non-fluorescent variant, rs-Eos3.1(H62G, Y63G), were included in each plate as controls. To prepare lysates for screening, frozen cell pellets were resuspended in 800 μl lysis buffer (50 mM Tris, 150 mM NaCl, pH 8) containing 50% BPER (Thermo Fisher) and shaken at 30°C for 30 min. Cell debris was pelleted by centrifugation and 95 μl of the cleared lysate was transferred to four separate 96-well microplates (Supplementary Fig. 2). Two of the plates were mixed with 5 μl calcium chloride solution to a final concentration of 0.5 mM. The other two plates were mixed with 5 μl EGTA solution to a final concentration of 1.0 mM. Green and red fluorescence intensities were measured on an Infinity M1000 fluorescence plate reader (Tecan). The plates were then illuminated with a 490 nm LED array (170 mW/cm^2^, Luxeon) for 15 s before reading fluorescence intensities again. One of the original plates in high calcium condition was swapped to high EGTA condition by adding 10 μl EGTA solution to a final concentration of 10 mM. Similarly, one of the original plates in high EGTA condition was swapped to high calcium condition by adding 10 μl calcium chloride solution to a final concentration of 5 mM. Fluorescence intensities were measured again. Finally, the plates were illuminated with a 405 nm LED array (200 mW/cm^2^, Loctite) for 10 s before reading final fluorescence intensities.

Library variants were selected based on four criteria: (1) contrast in green fluorescence following 490nm light illumination +/− Ca^2+^, quantified as:

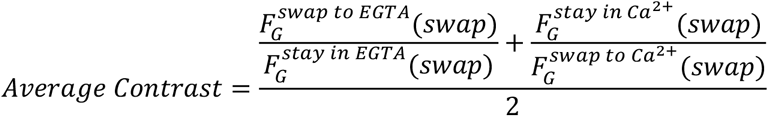

(2) recovery of fluorescence intensity following 400 nm light illumination, (3) minimum fluorescence change due to calcium binding in the absence of light illumination (indicator behavior), quantified as:

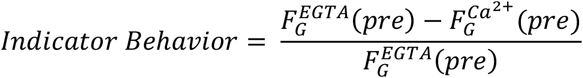

and (4) green brightness. We selected a total of 96 interesting library clones for sequencing, resulting in 19 unique sequences. These variants were expressed and purified to measure relative brightness, indicator behavior, and photoswitching kinetics in the presence and absence of Ca^2+^. One clone was renamed as rsCaMPARI (“rsCaMPARI-46” in Supplementary Table 1) and characterized further.

### rsCaMPARI expression and purification

rsCaMPARI protein was expressed with a N-terminal His_6_ tag in T7 express *E. coli* (New England Biolabs) using a pRSET expression vector. The bacteria were grown in auto-induction media using the Studier method^31^ for 36 h at 30°C. Cells were pelleted by centrifugation, lysed in Bacterial Protein Extraction Reagent (BPER)(Thermo Fisher), and the cleared lysate was loaded onto a column containing Ni-NTA agarose resin (QIAGEN). The column was washed with TBS (19.98 mM Tris, 136 mM NaCl, pH 7.4) + 10 mM imidazole before eluting with TBS + 200 mM imidazole. The eluted protein was loaded onto a Superdex 75 size-exclusion column (GE Healthcare) equilibrated with TBS and apparent monodispersity was confirmed. Fluorescent fractions were collected, combined, and stored at 4°C.

### In vitro analysis of purified protein

#### Absorbance and emission spectra

Absorption spectra of rsCaMPARI were measured in TBS with 0.5 mM CaCl_2_ or 1 mM EGTA on a NanoDrop One^c^ UV-VIS spectrophotometer (Thermo) in cuvettes with 10 mm path length. Emission spectra were measured on an Infinity M1000 plate reader (Tecan) set to 495 nm excitation and scanning emission from 505-700 nm.

#### Photophysical measurements

All the measurements were performed in 39 μM free calcium (+Ca) buffer (30 mM MOPS, 10 mM CaEGTA in 100 mM KCl, pH 7.2) or 0 μM free calcium (−Ca) buffer (30 mM MOPS, 10mM EGTA in 100mM KCl, pH 7.2). Absorbance measurements were performed using a UV-Vis spectrometer (Lamda 35, Perkin Elmer). Extinction coefficients were determined with the use of alkali denaturation method using extinction coefficient of denatured GFP as a reference (ε = 44000 M^−^ 1cm^−1^ at 447nm). Quantum Yield measurements were performed using an integration sphere spectrometer (Quantaurus, Hamamatsu) for proteins in +Ca buffer.

#### Two-photon measurements

The two-photon excitation spectra were performed as previously described^11^. Protein solution of 1-2 μM concentration in +Ca or −Ca buffer was prepared and measured using an inverted microscope (IX81, Olympus) equipped with a 60x/1.2 NA water immersion objective (Olympus). Two-photon excitation was obtained using an 80 MHz Ti-Sapphire laser (Chameleon Ultra II, Coherent) for spectra from 710 nm to 1080 nm. Fluorescence collected by the objective was passed through a short pass filter (720SP, Semrock) and a band pass filter (550BP88, Semrock), and detected by a fiber-coupled Avalanche Photodiode (APD) (SPCM_AQRH-14, Perkin Elmer). The obtained two-photon excitation spectra were normalized for 1 μM concentration and further used to obtain the action cross-section spectra (AXS) with fluorescein as a reference^32,33^.

#### Photoswitching rate measurements

To measure photoswitching rates +/− calcium, purified rsCaMPARI in low (10 mM EGTA) or high (10 mM CaEGTA) calcium conditions buffered with 25 mM Tris, 100 mM KCl, pH 7.5 was illuminated with a 470 nm LED (Mightex) with different bandpass filters (parts: FB450-10, FB460-10, FB470-10, FB480-10, FB490-10, and FB500-10, Thorlabs; ET485/25x, Chroma). Output spectra were measured with a USB4000-UV-VIS Ocean Optics spectrometer. Light intensities were measured on a power meter (Coherent #1098580) using a silicon photodiode (Coherent #1098313). Green fluorescence was measured on a plate reader (Tecan) at various timepoints and a single exponential rate was fitted (Prism, Graphpad) to the change in green fluorescence.

#### Determination of optimal marking and erasing light wavelengths for rsCaMPARI

We observed that the off-switching rate contrast was dependent on the wavelength of blue light used and that using lower wavelengths resulted in better off-switching rate contrast (~10-fold at 500 nm to ~25-fold at 450 nm) (Supplementary Fig. 7a-c). However, when exposed to wavelengths <470 nm, rsCaMPARI never reached the fully off-switched state and instead settled at an intermediate level of fluorescence (Supplementary Fig. 7b). We attributed this behavior to spectral overlap of the switching light with the broad 392 nm peak that drives rsCaMPARI on-switching. Indeed, when starting from the fully off-switched state, 450-470 nm light on-switches the protein (Supplementary Fig. 7d). Therefore, the observed intermediate resting fluorescence is due to 450-470 nm light driving both on- and off-switching to achieve an intermediate equilibrium state. Wavelengths between 470 nm and 490 nm should be used with rsCaMPARI to maintain high off-switching rate contrast while avoiding on-switching. We define the 470-490 nm region for rsCaMPARI as “marking light.” Likewise, we define the 390-410 nm region, which drives only on-switching, as “erasing light”.

#### K_d_ measurements

Solutions of EGTA-buffered free Ca^2+^ were prepared using a pH-titration method as previously described^34^ in 25 mM Tris, 100 mM KCl, pH 7.5. Free Ca^2+^ concentration was calculated using the following parameters for EGTA solutions with 0.1 M ionic strength at 25°C^35^: log *K*_Ca_ = 10.86, p*K*_1_ = 9.51, p*K*_2_ = 8.90. Note that p*K*_1_ and p*K*_2_ are adjusted 0.11 higher for 0.1 M ionic strength as explained by Tsien and Pozzan^34^. Therefore, the effective *K*_d_(Ca) for EGTA is calculated as follows:

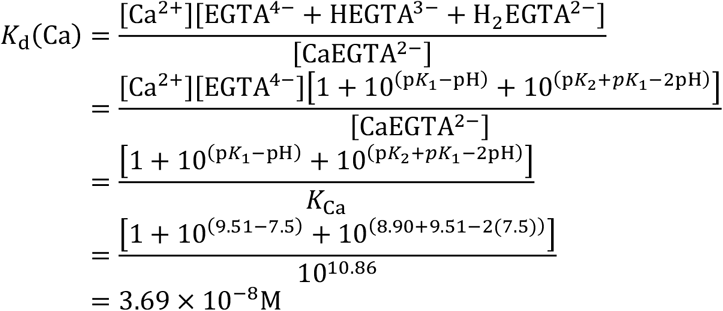

Ca-EGTA solutions over a range of free Ca^2+^ concentrations from 5.5 nM to 19 μM were prepared by mixing various volumes of a 10 mM Ca-EGTA solution with a 10 mM EGTA solution. To measure apparent affinity of rsCaMPARI for calcium ions, 2 μl of protein solution (~50 μM) was mixed with 98 μl of different Ca-EGTA solutions and the fluorescence intensity (Ex. 500 nm, Em. 515 nm) was measured on an Infinity M1000 fluorescence plate reader (Tecan). The data were fit (Prism, Graphpad) to a binding curve of the form:

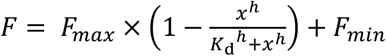

where *F* = fluorescence signal (AU)

*F*_*max*_ = maximum fluorescence signal (AU)

*F*_*min*_ = minimum fluorescence signal (AU)

*x* = concentration of free Ca^2+^ (M)

*K*_*d*_ = dissociation constant (M)

*h* = Hill coefficient

*F*_*max*_ was normalized to 1 to plot the relative fluorescence intensity. To measure rsCaMPARI off-switching rate as a function of free Ca^2+^ concentration, protein in Ca-EGTA solutions were prepared as described above and illuminated with various intervals of marking light (200 mW/cm^2^). The fluorescence intensity was measured at each timepoint to fit a single exponential.

The extrapolated kinetic rate *k* was then fitted to a binding curve of the form:

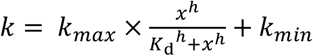

where *k* = kinetic rate (s^−1^)

*k*_*max*_ = maximum kinetic rate (s^−1^)

*K*_*min*_ = minimum kinetic rate (s^−1^)

*x* = concentration of free Ca^2+^ (M)

*K*_*d*_ = dissociation constant (M)

*h* = Hill coefficient

*k*_*min*_ was normalized to 1 to plot the relative off-switching rate.

### rsCaMPARI experiments in dissociated primary neuron cultures

All procedures involving animals were conducted in accordance with protocols approved by the Howard Hughes Medical Institute (HHMI) Janelia Research Campus Institutional Animal Care and Use Committee and Institutional Biosafety Committee. Hippocampal neurons extracted from P0 to 1 Sprague-Dawley rat pups were transfected with plasmids (see below) or plated directly without transfection into either 24-well glass-bottom plates (Mattek, #1.5 coverslip) for field electrode and channelrhodopsin experiments or 25 mm ultra-clean cover glasses (Sigma) for electrophysiology experiments. All glass surfaces were coated with poly-D-lysine prior to plating. Neurons were cultured in NbActiv4 medium (BrainBits) at 37°C with 5% CO_2_ in a humidified atmosphere. All imaging measurements were performed in imaging buffer containing 10 mM HEPES, 145 mM NaCl, 2.5 mM KCl, 10 mM glucose, 2 mM CaCl_2_, 1 mM MgCl_2_, pH 7.4 supplemented with synaptic blockers (10 μM CNQX, 10 μM CPP, 10 μM GABAZINE, and 1 mM MCPG; Tocris Bioscience) to block ionotropic glutamate, GABA, and metabotropic glutamate receptors^36^.

All images were acquired on an inverted Nikon Eclipse Ti2 microscope equipped with a SPECTRA X light engine (Lumencor), an AURA II light engine (Lumencor), and an ORCA-Flash 4.0 sCMOS camera (Hamamatsu). The SPECTRA X light engine was equipped with 485/25 and 395/25 nm excitation filters to produce marking and erasing light, respectively; and 550/15 and 640/30 nm excitation filters for exciting mRuby3 and JF635 dye, respectively. A quad bandpass filter (set number: 89000, Chroma) was used along with 525/36, 605/52, and 705/72 nm emission filters (Chroma) to image rsCaMPARI, mRuby3, and JF635 dye, respectively. A 550SP dichroic mirror (Thorlabs) was used to filter 560 nm light from the AURA II light engine for photostimulation of the ChrimsonR channelrhodopsin. Light powers were measured using a power meter (Thorlabs PM100A) with a Si photodiode (Thorlabs S120C or S170C). All image analysis and quantification were performed using Fiji software^37^.

### Characterization of rsCaMPARI in primary neuron cultures stimulated with a field electrode

Dissociated neurons were plated directly without transfection. 3 days after plating, the neurons were infected with AAV2/1 virus encoding rsCaMPARI-mRuby3 under control of the hsyn1 promoter. Imaging was performed 5-7 days after infection. The neurons were exposed to 10-20 s of continuous marking light (224 mW/cm^2^) through a 10x/0.45 NA objective (Nikon) for simultaneous photoswitching and imaging (10 Hz acquisition) of rsCaMPARI during one marking cycle. Each marking cycle was erased with 3 s of erasing light (224 mW/cm^2^ power). Field stimulations (80 Hz, 1 ms) were produced using a custom-built field electrode controlled by a high current isolator (A385, World Precision Instruments) set to 90 mA. Stimulations were controlled using an Arduino Uno board and synchronized with light sources in Nikon Elements software.

For characterizing rsCaMPARI spontaneous recovery in the dark, neurons that received prior exposure to 20 s of marking light +/− field stimulation from one marking cycle were incubated in the dark in a stage top incubator (Tokai Hit) set to 37°C. Snapshots of rsCaMPARI were acquired every 5 min (224 mW/cm^2^, 100 ms). In between each snapshot, 160 stimulations (80 Hz, 1ms) were delivered using the field electrode. After 60 min, rsCaMPARI was erased with ~3 s of erasing light (405 nm, 224 mW/cm^2^ power) before a final snapshot of rsCaMPARI was acquired. For characterizing rsCaMPARI photofatigue across multiple marking cycles, neurons were continually marked and erased with 20 s of marking light (224 mW/cm^2^) and 3 s of erasing light (224 mW/cm^2^ power), respectively. Field stimulations (3× 160 stims, 80 Hz, 1 ms) were delivered during marking light illumination on every odd cycle.

### Simultaneous electrophysiology and fluorescence imaging in primary neuron culture

Dissociated neurons were transfected with a pAAV-plasmid containing rsCaMPARI-mRuby3 under control of the hsyn1 promoter by electroporation (Lonza, P3 Primary Cell 4D-Nucleofector X kit) according to the manufacturer’s instruction. 9 days after plating, the neurons were exchanged to imaging buffer containing synaptic blockers for imaging. Within a field of view containing multiple neurons expressing rsCaMPARI, a single neuron was patched and the entire field of view was exposed to 15 s of marking light (150 mW/cm^2^) through a 40x/1.3 NA oil objective (Nikon) for simultaneous photoswitching and imaging (10 Hz acquisition) of rsCaMPARI during one marking cycle. Each marking cycle was erased with ~3 s of erasing light (486 mW/cm^2^).

Whole-cell patch clamp recordings were performed with filamented glass micropipettes (Sutter instruments) pulled to a tip resistance of 10-12 MΩ. The internal solution in the pipette contained (in mM): 130 potassium methanesulfonate, 10 HEPES, 5 NaCl, 1 MgCl2, 1 Mg-ATP, 0.4 Na-GTP, 14 Tris-phosphocreatine, adjusted to pH 7.3 with KOH, and adjusted to 300 mOsm with sucrose. Pipettes were positioned using a MPC200 manipulator (Sutter instruments) and current clamp traces were recorded using an EPC800 amplifier (HEKA) and digitized using a National Instruments PCIe-6353 acquisition board. To generate action potentials, current was injected to induce spike trains (3x 20-200 pA for 1 s) and voltage was monitored. WaveSurfer software (https://wavesurfer.janelia.org/) was used to control the amplifier, camera, light source, and record voltage and current traces.

### rsCaMPARI experiments in primary neurons with a subset driven by a channelrhodopsin

Dissociated neurons were transfected with a pAAV-plasmid containing a ChrimsonR and HaloTag^38^ fusion protein under control of the hsyn1 promoter by electroporation (Lonza, P3 Primary Cell 4D-Nucleofector X kit) according to the manufacturer’s instruction and mixed 2:1 with non-transfected cells before plating. 6 days after plating, the neurons were infected with AAV2/1 virus encoding rsCaMPARI-mRuby3 under control of hsyn1. Imaging was performed 7 days after infection. To label neurons expressing ChrimsonR-HaloTag, cultures were incubated with 100 nM JF635-HaloTag ligand for 30 min. Neurons were then washed in imaging buffer three times and the buffer was then replaced with imaging buffer containing synaptic blockers. The neurons were exposed to 10 s of continuous marking light (285 mW/cm^2^) through a 10x/0.45 NA objective (Nikon) for simultaneous photoswitching and imaging (10 Hz acquisition) of rsCaMPARI during one marking cycle. 560 nm light (59 mW/cm^2^) was also pulsed (10 ms pulses at 10 Hz) through the objective during the entire marking light illumination to fully drive the channelrhodopsin. Each marking cycle was erased with ~3 s of erasing light (153 mW/cm^2^ power). Classification of neurons expressing channelrhodopsin was done using a threshold of the JF635 fluorescence (Fiji).

### rsCaMPARI experiments in larval zebrafish (Danio rerio)

All zebrafish experiments were conducted in accordance with the animal research guidelines from the National Institutes of Health and were approved by the Institutional Animal Care and Use Committee and Institutional Biosafety Committee of Janelia Research Campus.

To generate the Tg[*elavl3*:rsCaMPARI-mRuby3]^jf93^ line, 1-2 cell embryos from *casper* background zebrafish were injected with a Tol2 vector containing rsCaMPARI-mRuby3 under control of the *elavl3* pan-neuronal promoter. Potential founders were screened for bright green and red fluorescence in the brain and later crossed with *casper* background zebrafish to screen for progeny (F1 generation) exhibiting pan-neuronal rsCaMPARI-mRuby3 expression in the central nervous system (CNS). The F1 generation fish were later incrossed and 4-5 days post-fertilization (dpf) larvae exhibiting bright green fluorescence in the CNS were used for experiments.

We observed poor correlation between the rsCaMPARI green signal and the mRuby3 red signal in the larvae (Supplementary Fig. 15), presumably because the larval zebrafish brain is rapidly developing and the maturation time of the mRuby3 chromophore is slow^39^ compared to rsCaMPARI. Therefore, we caution against using mRuby3 as an expression normalization tag in larval zebrafish. However, the mRuby3 tag was useful for initially locating and positioning the zebrafish brain within the field of view for imaging without using blue light, which would induce further rsCaMPARI photoswitching.

Zebrafish in system water were illuminated for 10 s with marking light (470 nm Mightex LED fitted with a Chroma 485/25x filter, 400 mW/cm^2^) under various stimulus conditions, including freely swimming in system water, anesthetized in 0.24 mg/mL tricaine methanesulfonate (MS-222, Sigma), cold water (4°C), warm water (45°C), and following 15-30 min exposure to 800 μM 4-aminopyridine (4-AP, Sigma). Following marking light exposure, the fish was anesthetized in 0.24 mg/mL MS-222 and immobilized with 2% agarose in a glass capillary tube (size 2, Zeiss) for imaging. For fish undergoing multiple cycles of marking and erasing, the fish was transferred to fresh system water and carefully removed from the agarose. The fish was allowed to recover to freely swimming behavior (typically within 15 min) and a brief 3 s illumination with erasing light (405 nm Loctite LED, 200 mW/cm^2^) was used to reset the sensor before another marking cycle began.

Larval zebrafish brains were imaged on a Zeiss Lightsheet Z.1 microscope equipped with 50 mW 405, 488, and 561 nm laser lines and a pco.edge 5.5 sCMOS camera (PCO). Images were acquired using either a 10x/0.5 NA (for whole brain) or 20x/1.0 NA (for forebrain) water immersion objective (Zeiss) for detection, and two 10x/0.2 NA optics (Zeiss) for dual-side illumination. The sample chamber was filled with system water containing 0.24 mg/mL MS-222 and the solidified agarose was partially extruded from the capillary tube to position the fish within the sample chamber. A 585 nm longpass emission filter was used to visualize mRuby3 fluorescence for positioning and a 505-545 nm bandpass emission filter was used to image rsCaMPARI fluorescence. Each Z slice was acquired in dual-side illumination mode with pivot scan using 1% 488 nm excitation and 100 ms exposure time. Two Z stacks (3 μm steps for whole brain or 2 μm steps for forebrain) were acquired using continuous drive mode in the following order: (1) acquire first stack as the “marked” dataset (green), (2) erase the stack with 5% 405 nm light, and (3) acquire second stack as the “reference” dataset (pseudo-colored magenta). Total time to acquire both marked and reference stacks was ~1-2 minutes.

Acquired images were dual-side fused in Zeiss ZEN software. Motion drift between marked and reference stacks were corrected using affine alignment tools in Computational Morphometry ToolKit^40^ (CMTK) software (https://www.nitrc.org/projects/cmtk/) with the following parameters: exploration 26, accuracy 0.4, and dofs 12. Composite marked and reference images and ΔF/F images were created in Fiji^37^. Here, ΔF/F is defined as:

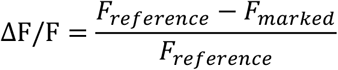

To compute mean correlation coefficients between trials in the same fish, reference image stacks of the forebrain were first aligned to each other with an initial affine alignment in CMTK with the following parameters: exploration 26, accuracy 0.4, and dofs 12; the images were then further aligned with a restrained warp in CMTK with the following parameters: exploration 26, coarsest 4, grid-spacing 90, refine 3, accuracy 0.4, and jacobian-weight 0.05. Marked image stacks were reformatted in CMTK to align with its respective reference stack and ΔF/F images were calculated. The resulting ΔF/F image stacks were cropped to remove sub-pallium regions where excitation light was scattered by pigmentation in the eyes of the zebrafish, and activated ensembles were selected by a 0.25 ΔF/F threshold. Representative slices showing selected activated ensembles are shown in Supplementary Fig. 16. Mean correlations between activated ensemble image stacks were calculated using cosine similarity across all slices using a custom macro in Fiji according to the following formula:

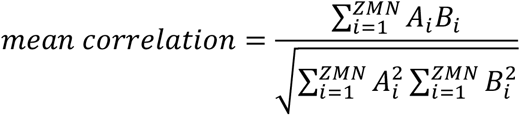

where A and B are image stacks with Z slices, and each slice has dimension M × N pixels.

