## Supplemental Figures and Table for "rsCaMPARI: an erasable marker of neuronal activity"

**-Supplementary Figures 1-16**
**-Supplementary Table 1**

### Supplementary Figures

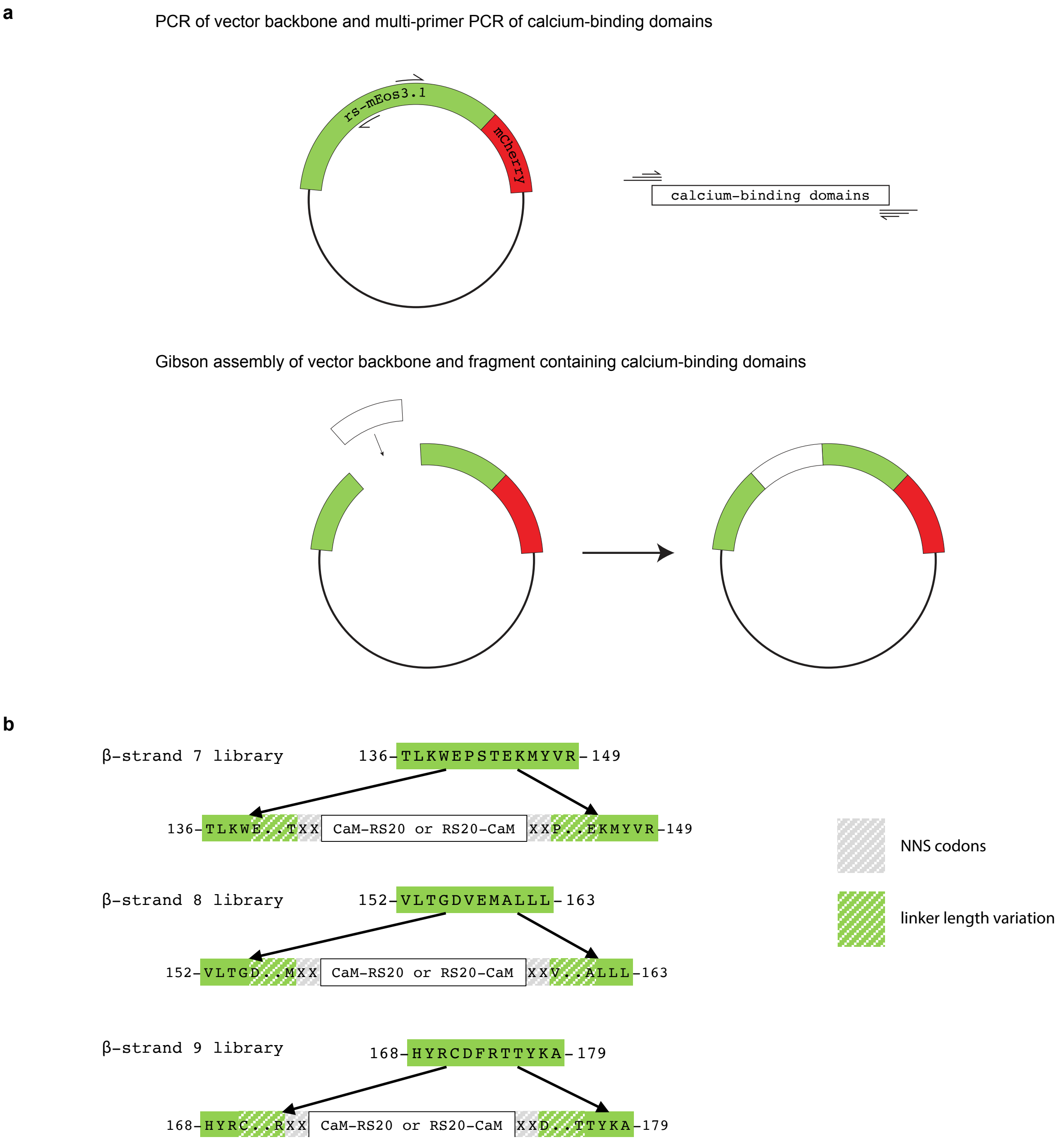


**Supplementary Figure 1: Design and construction of libraries to engineer an erasable calcium activity marker. a**, Outline of Gibson assembly strategy for constructing insertion libraries of calmodulin-binding domains into a reversibly photoswitchable variant of mEos3.1 (rs-mEos3.1). **b**, Theoretical amino acid composition of insertion variants for each β-strand library. Partially shaded grey boxes are randomized amino acids encoded by NNS codons and partial shaded green boxes are linker length variants from 0 to 4 amino acids extended from the native β-strand. For example, the N-terminal linker of the β-strand 7 library could be XX, EXX, EPXX, EPSXX, or EPSTXX. Similarly, the C-terminal linker could be XX, XXE, XXTE, XXSTE, or XXPSTE. Theoretical diversity for each β-strand library is 8 x $10^6$.

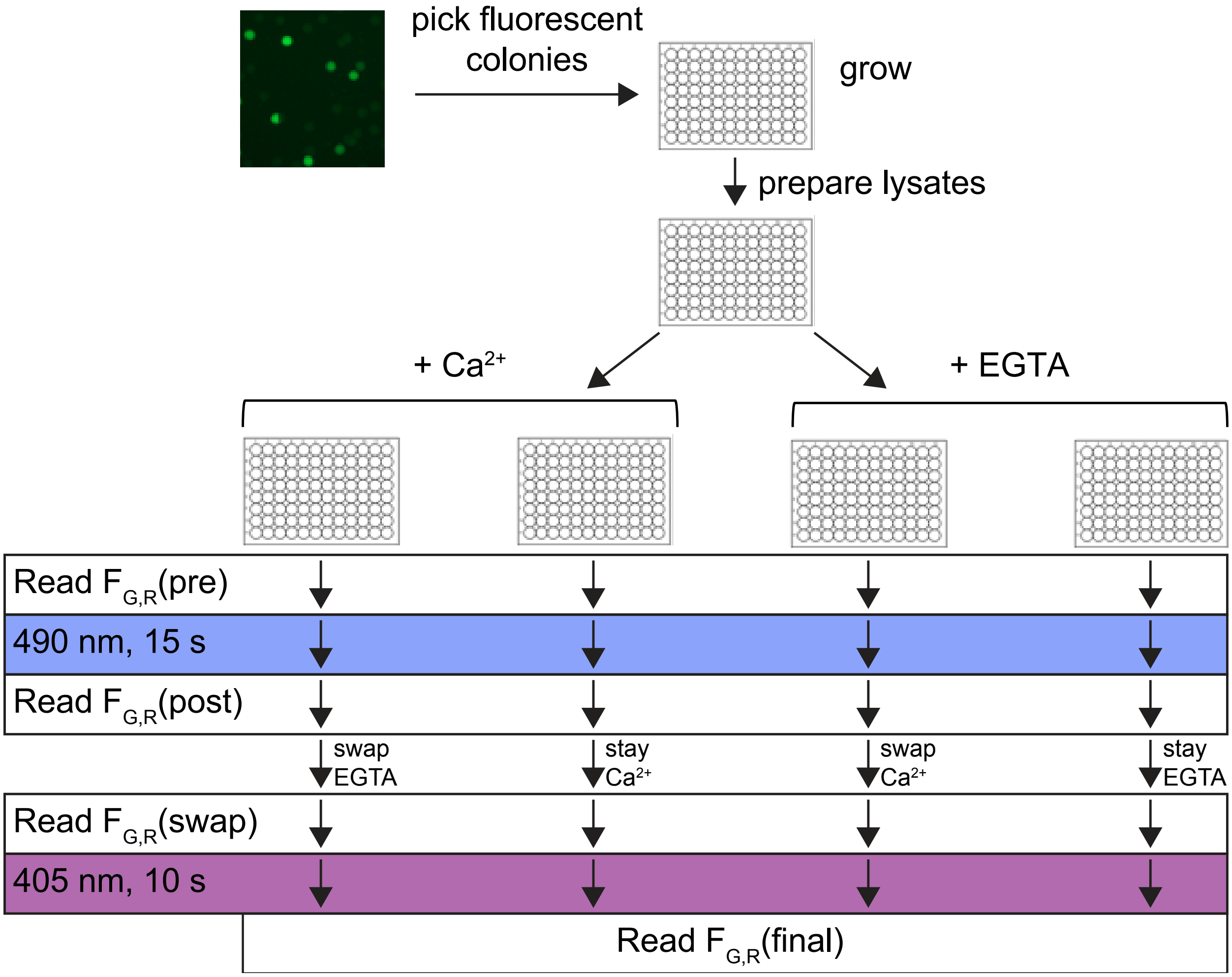


**Supplementary Figure 2: Outline of library screening to engineer rsCaMPARI.** Light intensities for 490 nm and 405 nm light illumination are 170 mW/cm$^2$ and 200 mW/cm$^2$, respectively. Initial concentration for $Ca^{2+}$ and EGTA are 0.5 mM and 1 mM, respectively. Concentrations for swapped $Ca^{2+}$ and EGTA are 5 mM and 10 mM, respectively.

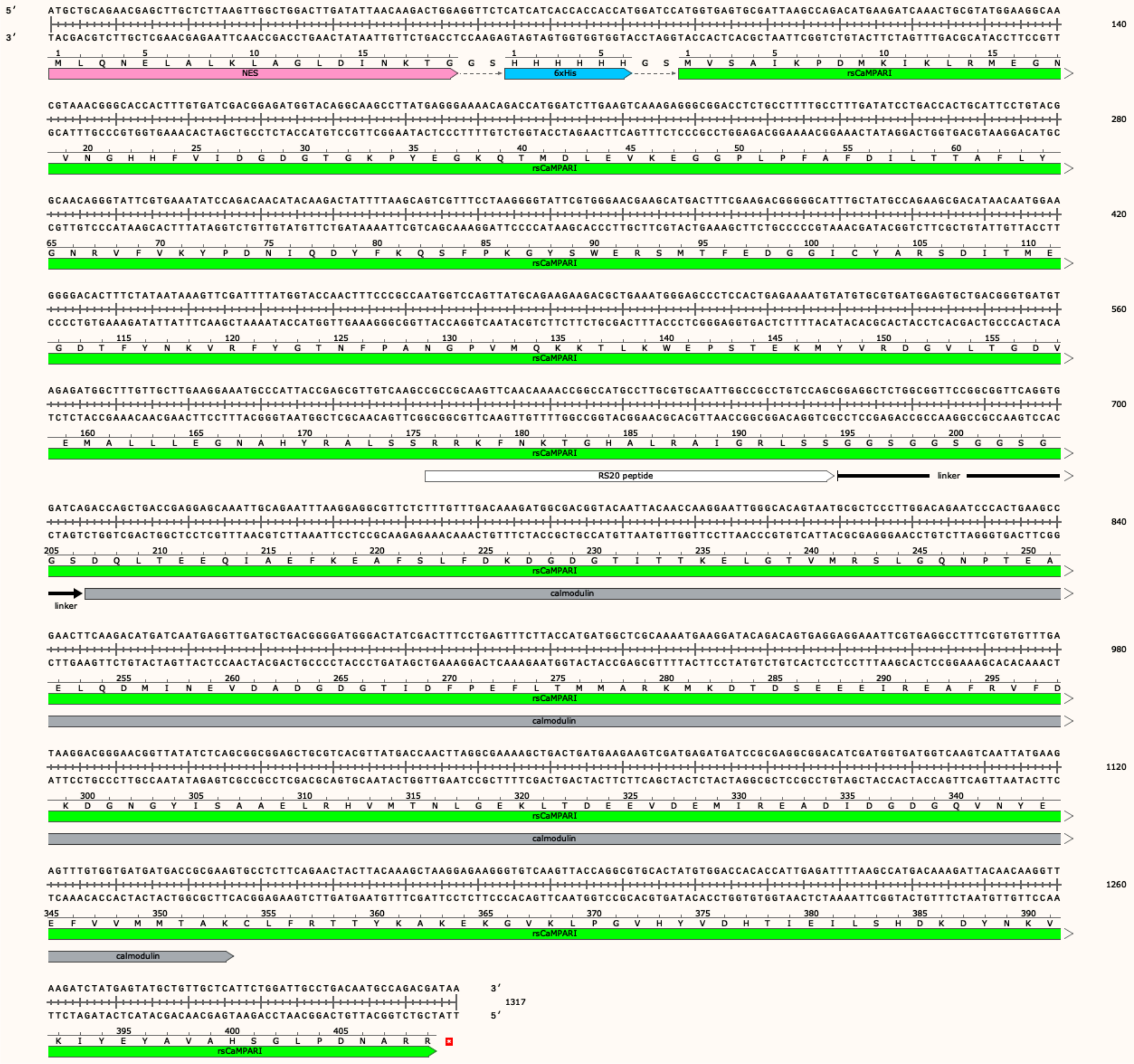


**Supplementary Figure 3: DNA and amino acid sequence of rsCaMPARI with sequence features annotated.** A nuclear export signal (NES) is added to the N-terminus for excluding rsCaMPARI from the nucleus when expressed in eukaryotic cells.

```
mEos3.1        M-SAIKPDMKIKLRMEGNVNGHHFVIDGDGTGKPFEGKQSMDLEVKEGGPLPFAFDILTT  59
rsCaMPARI      MVSAIKPDMKIKLRMEGNVNGHHFVIDGDGTGKPYEGKQTMDLEVKEGGPLPFAFDILTT  60

mEos3.1        AFHYGNRVFAKYPDNIQDYFKQSFPKGYSWERSLTFEDGGICNARNDITMEGDTFYNKVR  119
rsCaMPARI      AFLYGNRVFVKYPDNIQDYFKQSFPKGYSWERSMTFEDGGICYARSDITMEGDTFYNKVR  120

mEos3.1        FYGTNFPANGPVMQKKTLKWEPSTEKMYVRDGVLTGDVEMALLLEGNAHYR---------  170
rsCaMPARI      FYGTNFPANGPVMQKKTLKWEPSTEKMYVRDGVLTGDVEMALLLEGNAHYRALSSRRKFN  180

mEos3.1        ------------------------------------------------------------  170
rsCaMPARI      KTGHALRAIGRLSSGGSGGSGGSGGSDQLTEEQIAEFKEAFSLFDKDGDGTITTKELGTV  240

mEos3.1        ------------------------------------------------------------  170
rsCaMPARI      MRSLGQNPTEAELQDMINEVDADGDGTIDFPEFLTMMARKMKDTDSEEEIREAFRVFDKD  300

mEos3.1        -----------------------------------------------------CDFRTTY  177
rsCaMPARI      GNGYISAAELRHVMTNLGEKLTDEEVDEMIREADIDGDGQVNYEEFVVMMTAKCLFRTTY  360

mEos3.1        KAKEKGVKLPGAHFVDHCIEILSHDKDYNKVKLYEHAVAHSGLPDNARR    226
rsCaMPARI      KAKEKGVKLPGVHYVDHTIEILSHDKDYNKVKIYEYAVAHSGLPDNARR    409
```

**Supplementary Figure 4: Amino acid sequence comparison of rsCaMPARI with mEos3.1.** Point mutations between rsCaMPARI and mEos3.1 outside of the calcium-binding domains are highlighted in yellow. The chromophore is underlined, the RS20 calmodulin-binding peptide is outlined with a box, and calmodulin is shaded in grey.

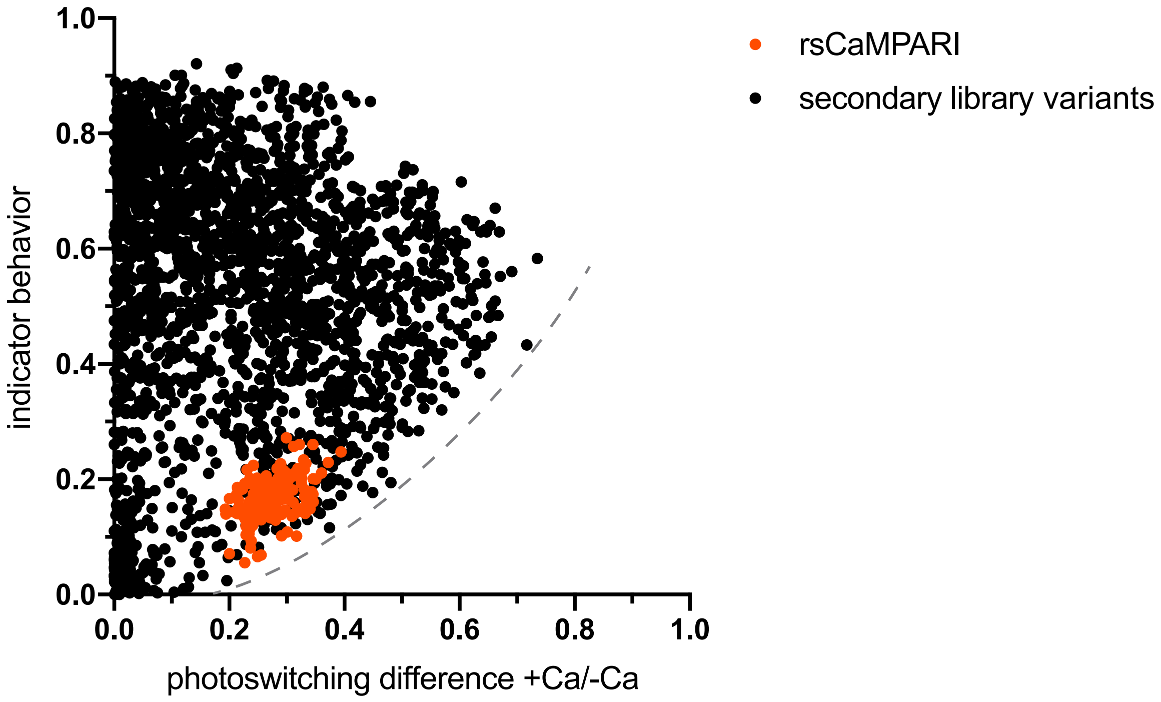


**Supplementary Figure 5: rsCaMPARI secondary library screen showing relationship between photoswitching contrast and indicator behavior.** rsCaMPARI controls are shown in red and secondary library variants are shown in black. A dashed grey line illustrates a boundary beyond which no variants with high photoswitching contrast and low indicator behavior were observed.

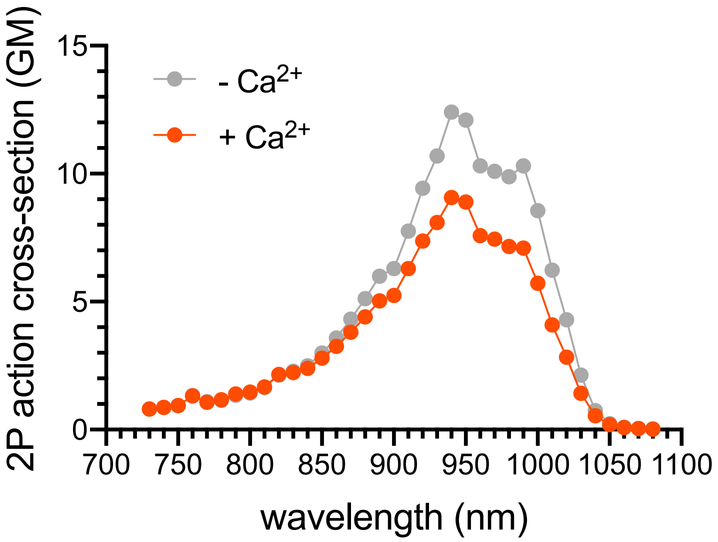


**Supplementary Figure 6: Two-photon action cross-section of rsCaMPARI in the presence and absence of $Ca^{2+}$.**

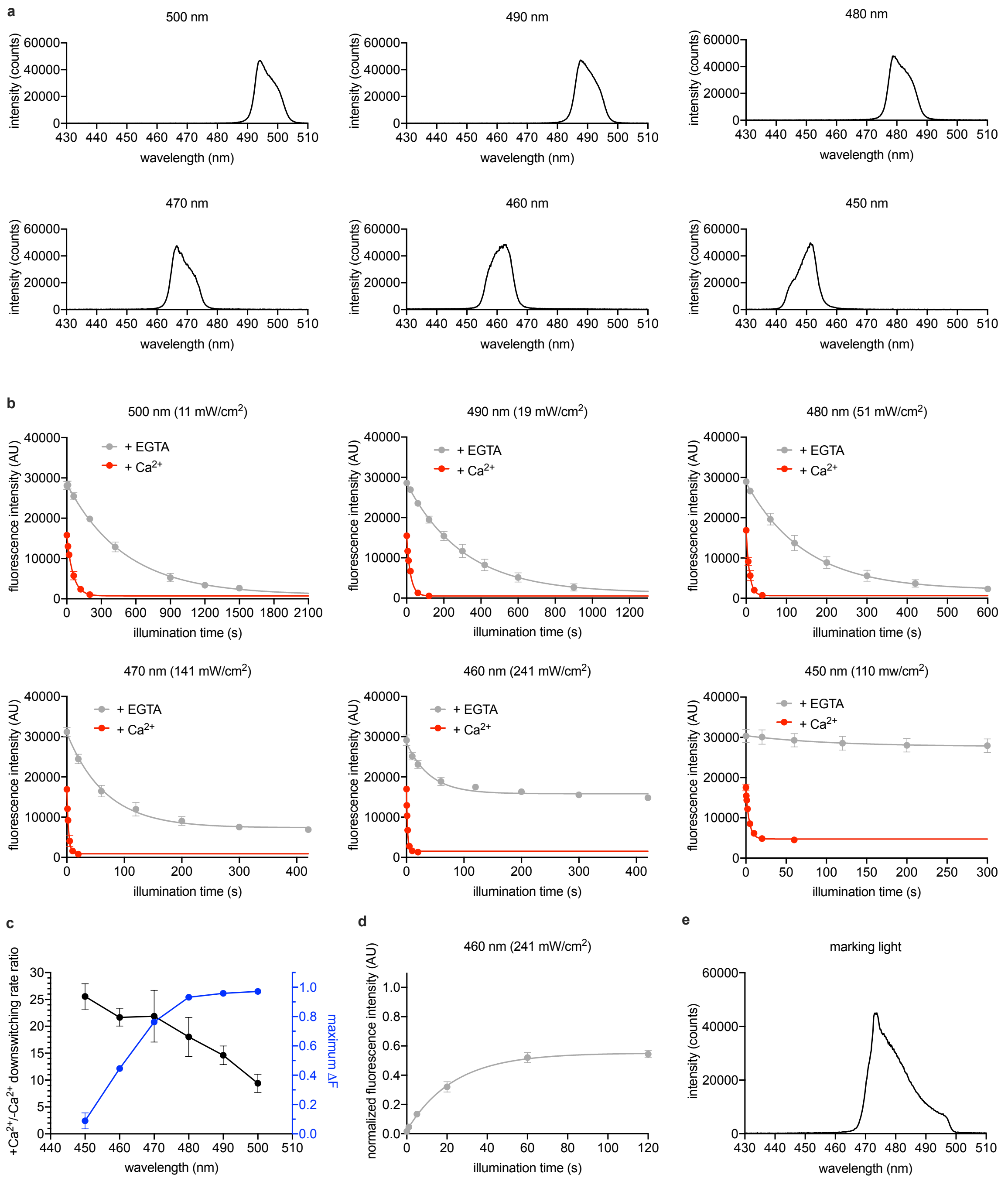


**Supplementary Figure 7: Optimization of blue light wavelengths for off-switching of rsCaMPARI. a**, Spectra of bandpass filters tested. **b**, Off-switching time-course of rsCaMPARI in the presence or absence of calcium under illumination of blue light with spectra shown in (A). Lines are single-exponential fits to data. Error bars are standard deviation, n=4 replicate measurements. **c**, Off-switching rate contrast and maximum fluorescence change (ΔF) as a function of wavelength. Error bars are standard deviation, n=4 replicate measurements. **d**, On-switching time-course of rsCaMPARI from dim off-state under illumination with 460 nm light. Line is single-exponential fit to data. Error bars are standard deviation, n=4 replicate measurements. **e**, Spectra of marking light from a Mightex 470 nm LED filtered through a Chroma 485/25x filter.

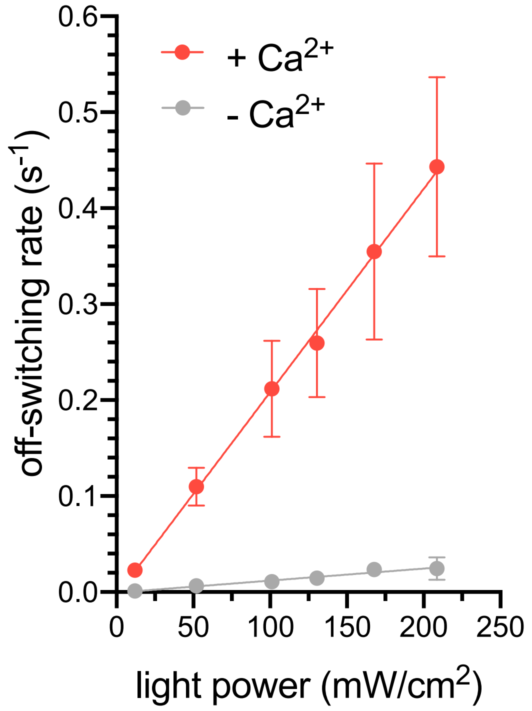


**Supplementary Figure 8: The relationship between rsCaMPARI off-switching rate and light power intensity.** Fitted lines are a simple linear regression to data. Error bars are standard deviation, n=4 replicate measurements.

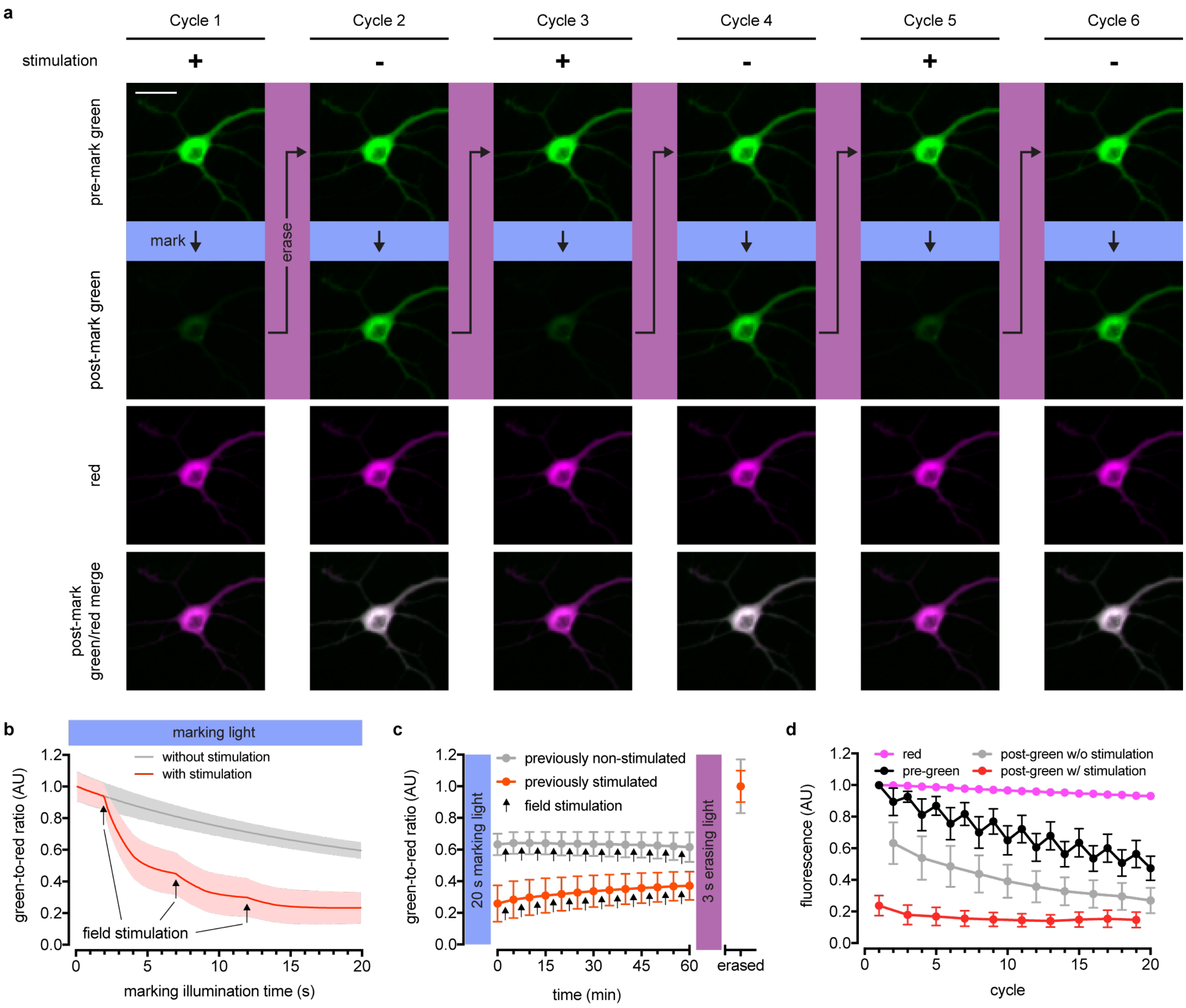


**Supplementary Figure 9: rsCaMPARI marks neurons stimulated by a field electrode. a**, Fluorescence images of rsCaMPARI-mRuby3 in a representative neuron undergoing multiple cycles of exposure to a 10 s window of marking light (224 mW/cm$^2$) +/- stimulation (2x 160 stims at 80 Hz). Each cycle was reset with a 3 s pulse of erasing light (224 mW/cm$^2$). Scale bar is 30 μm. **b**, Fluorescence time-course traces of neurons undergoing one cycle of illumination +/- stimulation (3x 160 stims at 80 Hz). Arrows on time-course trace denote start of each stimulation. Error bars are standard deviation, n=15 neurons from two independent wells. **c**, Time-course of rsCaMPARI spontaneous recovery in the dark at 37°C following marking light illumination of previously non-stimulated or previously stimulated neurons. Arrows denote a bout of field stimulation between each imaging timepoint (160 stims at 80 Hz). After one hour, the neurons were reset with a 3 s pulse of erasing light. Error bars are standard deviation, previously non-stimulated n=66 neurons and previously stimulated n=59 neurons from two independent wells. **d**, Photofatigue of rsCaMPARI over successive cycles of marking light illumination with or without field stimulation. Each cycle is followed by erasing light to reset the sensor. Error bars are standard deviation, n=7 neurons from three independent wells.

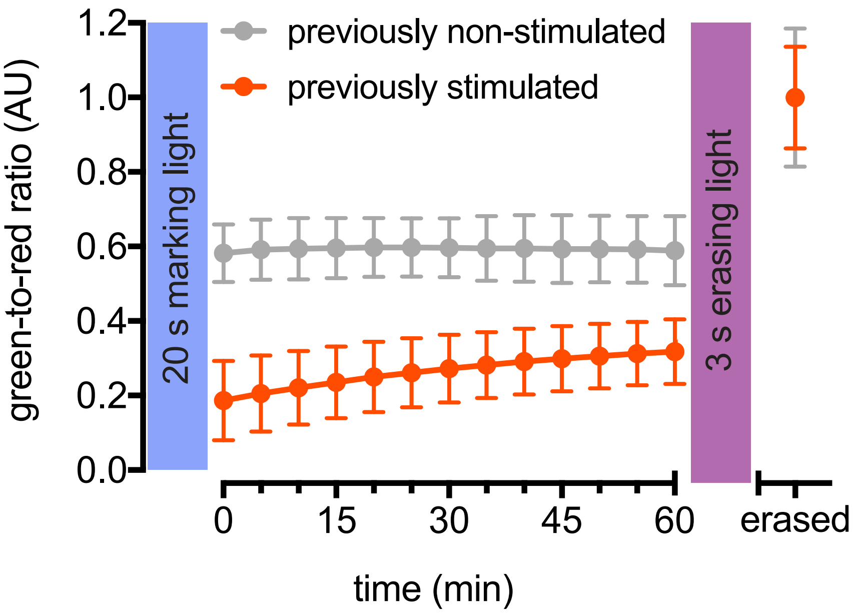


**Supplementary Figure 10: Spontaneous recovery of rsCaMPARI.** Time-course of rsCaMPARI spontaneous recovery in the dark at 37°C following blue light illumination of previously non-stimulated or previously stimulated neurons. No stimulations were delivered between each imaging timepoint. After one hour, the neurons were reset with a 3 s pulse of erasing light. Error bars are standard deviation, previously non-stimulated n=66 neurons and previously stimulated n=66 neurons from two independent wells.

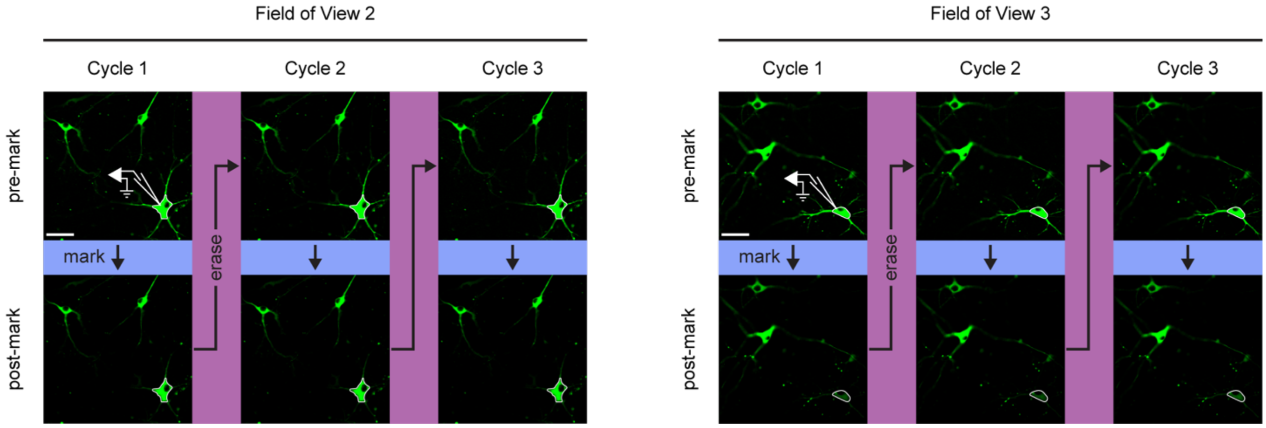


**Supplementary Figure 11: rsCaMPARI selectively marks neurons stimulated by current injection through a patch pipette.** Fluorescence images of rsCaMPARI expressed in dissociated primary rat hippocampal neurons before and after 15 s of marking light illumination. A single cell, denoted by pipette drawing, is patched and stimulated during marking light illumination. Scale bar is 50 μm. Quantification of $F_{pre}/F_{post}$ ratios are shown in Fig. 2G.

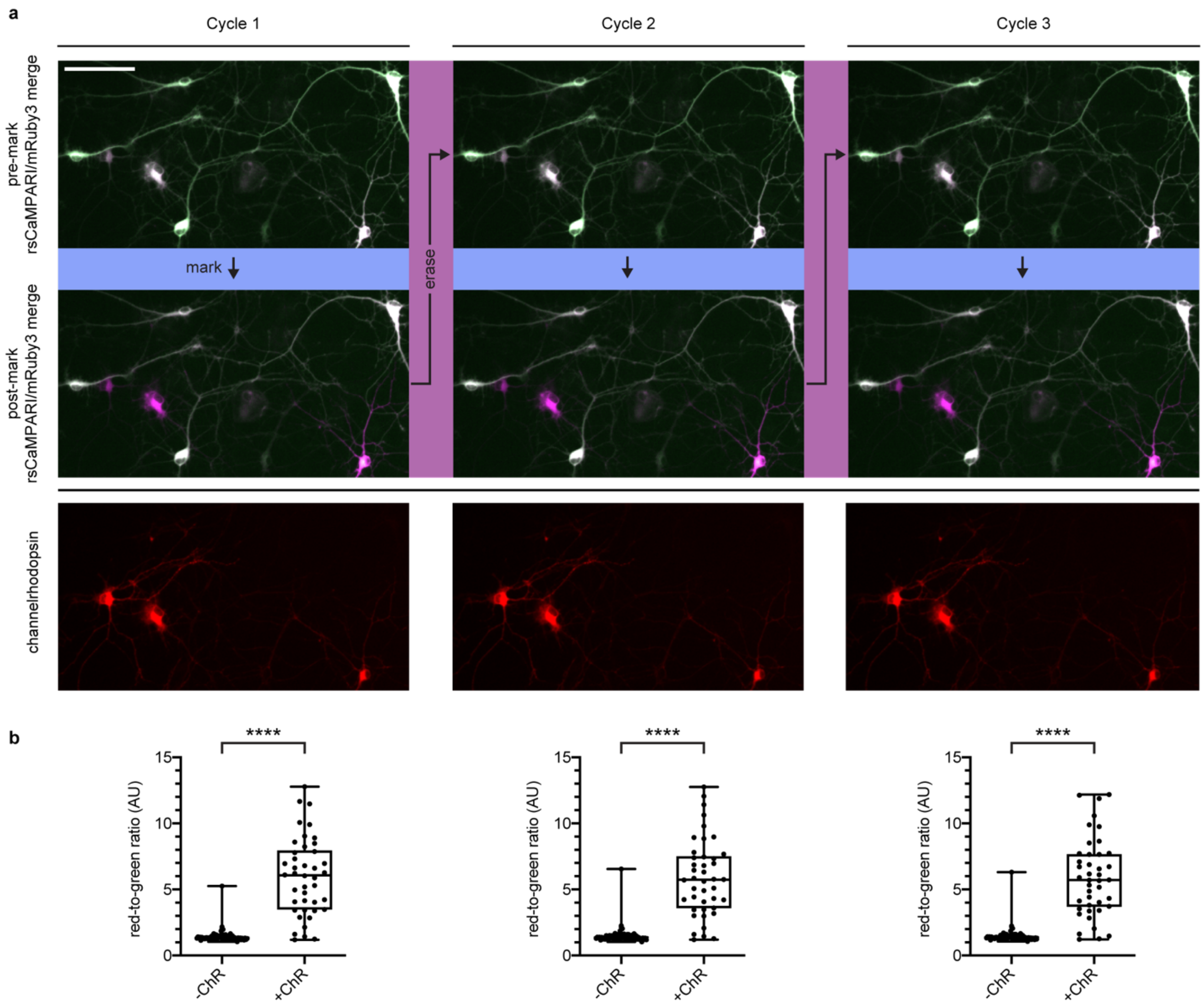


**Supplementary Figure 12: rsCaMPARI selectively marks neurons activated by a channelrhodopsin. a**, Merged rsCaMPARI (green) and mRuby3 (magenta) fluorescence images pre and post marking light illumination (285 mW/cm$^2$, 10 s) from three cycles. Neurons expressing ChrimsonR-HaloTag labeled with JF635 HaloTag ligand (red) are shown in bottom panels. Scale bar is 100 μm. **b**, Relative red-to-green fluorescence ratios of -ChR and -+ChR neurons post-marking light illumination. +ChR n=42 neurons and -ChR n=79 neurons from 17 independent wells. ****P < 0.0001, Wilcoxon rank-sum test.

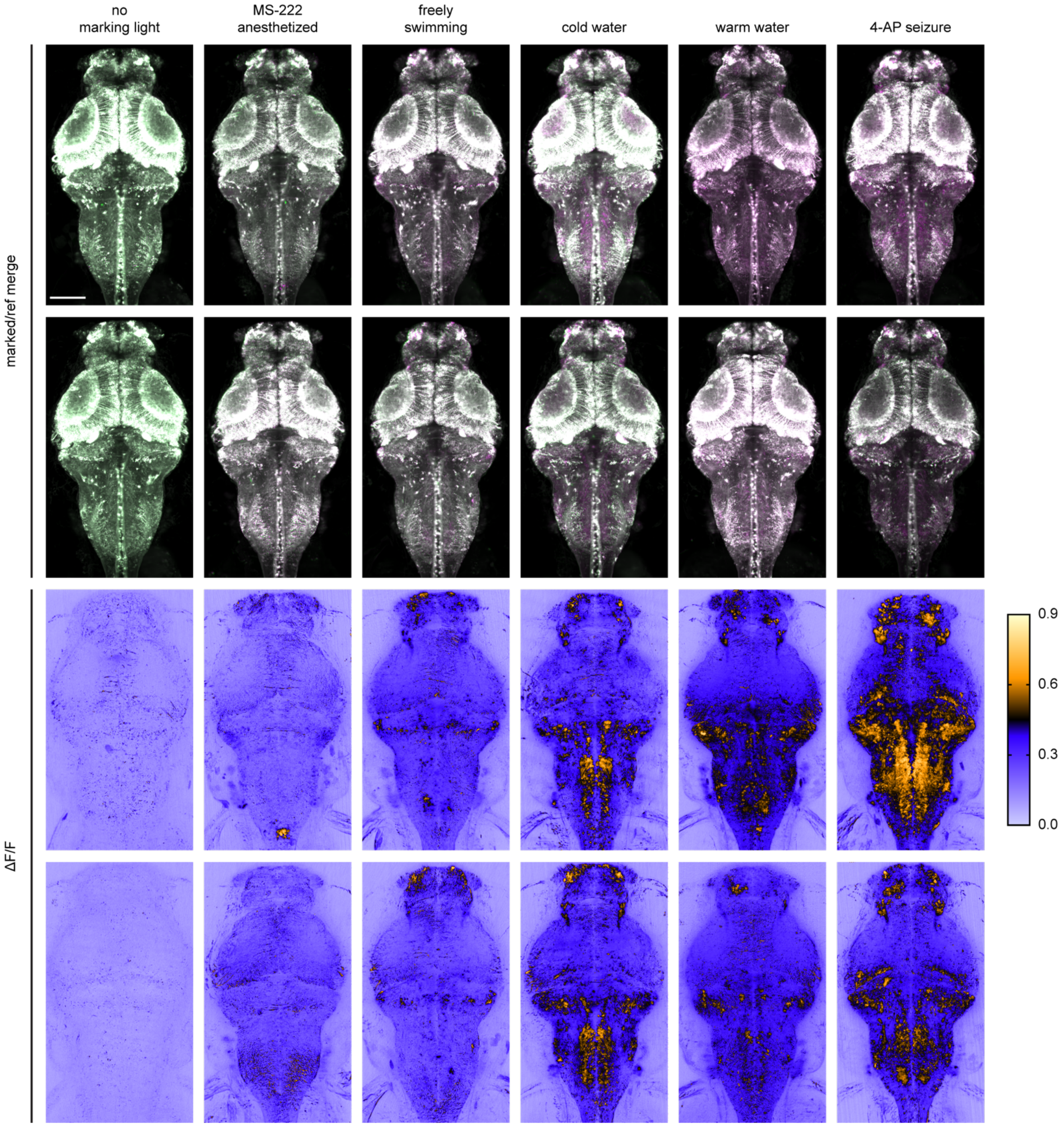


**Supplementary Figure 13: Replicates of rsCaMPARI labeling in zebrafish brain after exposure to different stimuli.** Each image is a maximum intensity Z projection of the entire brain from zebrafish larvae (4 to 5 dpf). Top half of images are merged reference and erased images, pseudo-colored green and magenta, respectively. Bottom half of images are corresponding ΔF/F images. Imaging conditions and brightness/contrast are identical to images shown in Fig. 3C. Scale bar is 100 μm.

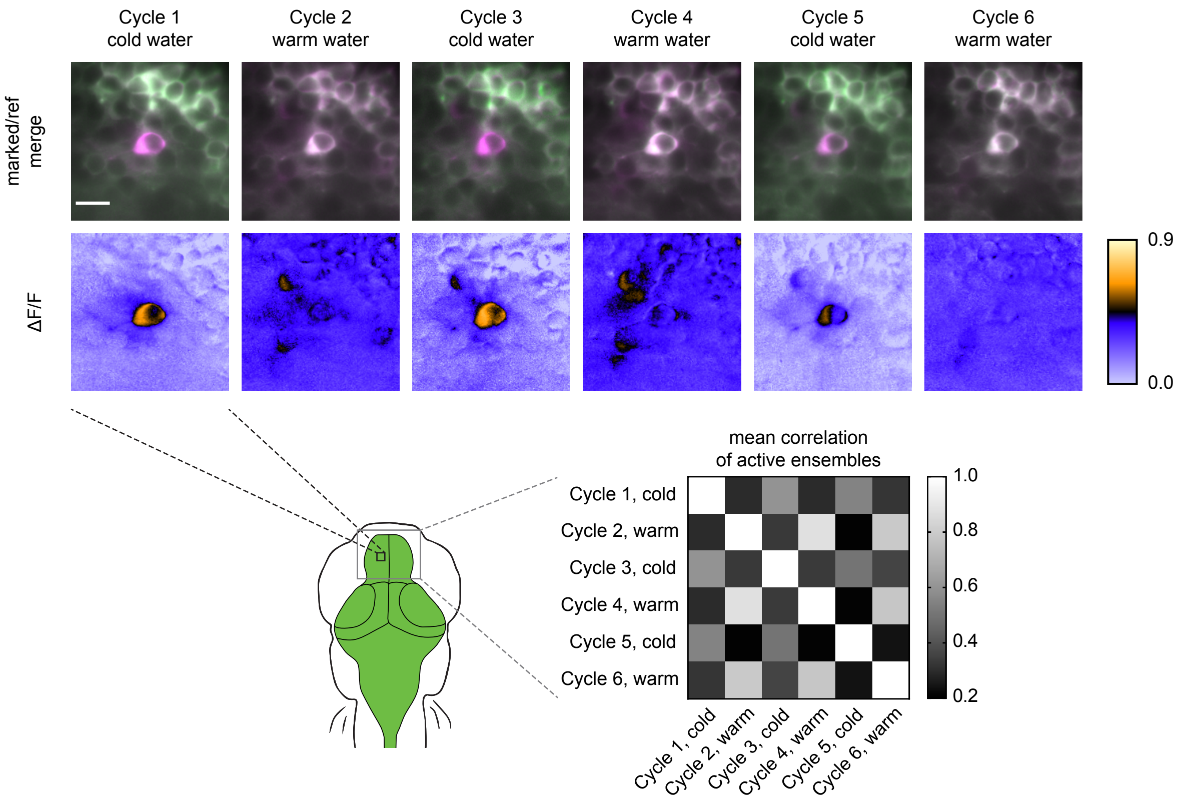


**Supplementary Figure 14: Replicate of multiple rsCaMPARI labeling cycles in the same zebrafish.** Top panels are individual Z slices from the pallium of the same fish (5 dpf) brain illustrating the same field of view from 6 marking cycles of alternating cold and warm water stimulus. Scale bar is 10 μm. Bottom-right panel is a normalized correlation matrix comparing labeling patterns in the upper pallium and habenula across multiple marking cycles in the same fish.

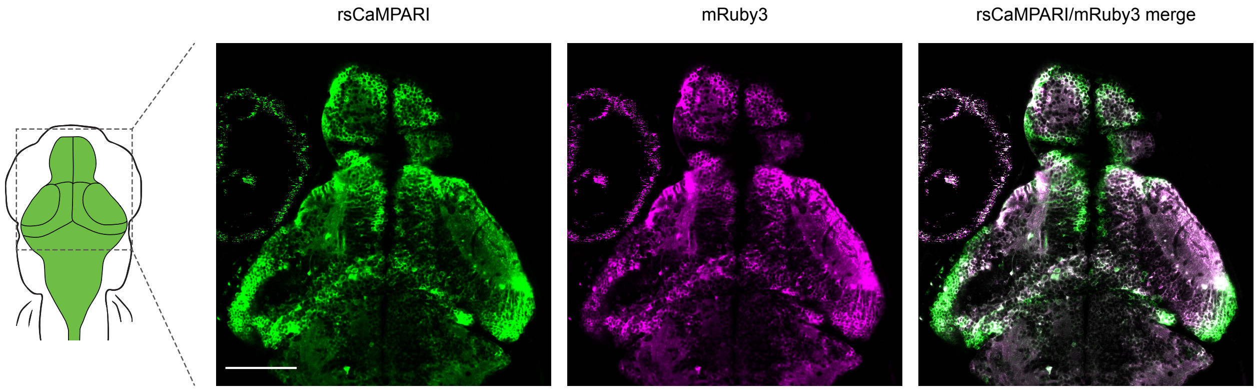


**Supplementary Figure 15: mRuby3 is poor for normalizing rsCaMPARI expression in larval zebrafish.** Two-photon image of rsCaMPARI-mRuby3 in 5 dpf zebrafish brain without prior exposure to marking light. Excitation wavelength is 960 nm with detection wavelength ranges of 490-534 nm and 570-695 nm for rsCaMPARI and mRuby3, respectively. Note the general lack of mRuby3 fluorescence in nascent neurons in the periphery of brain structures, presumably due to its long maturation time. Scale bar is 100 μm.

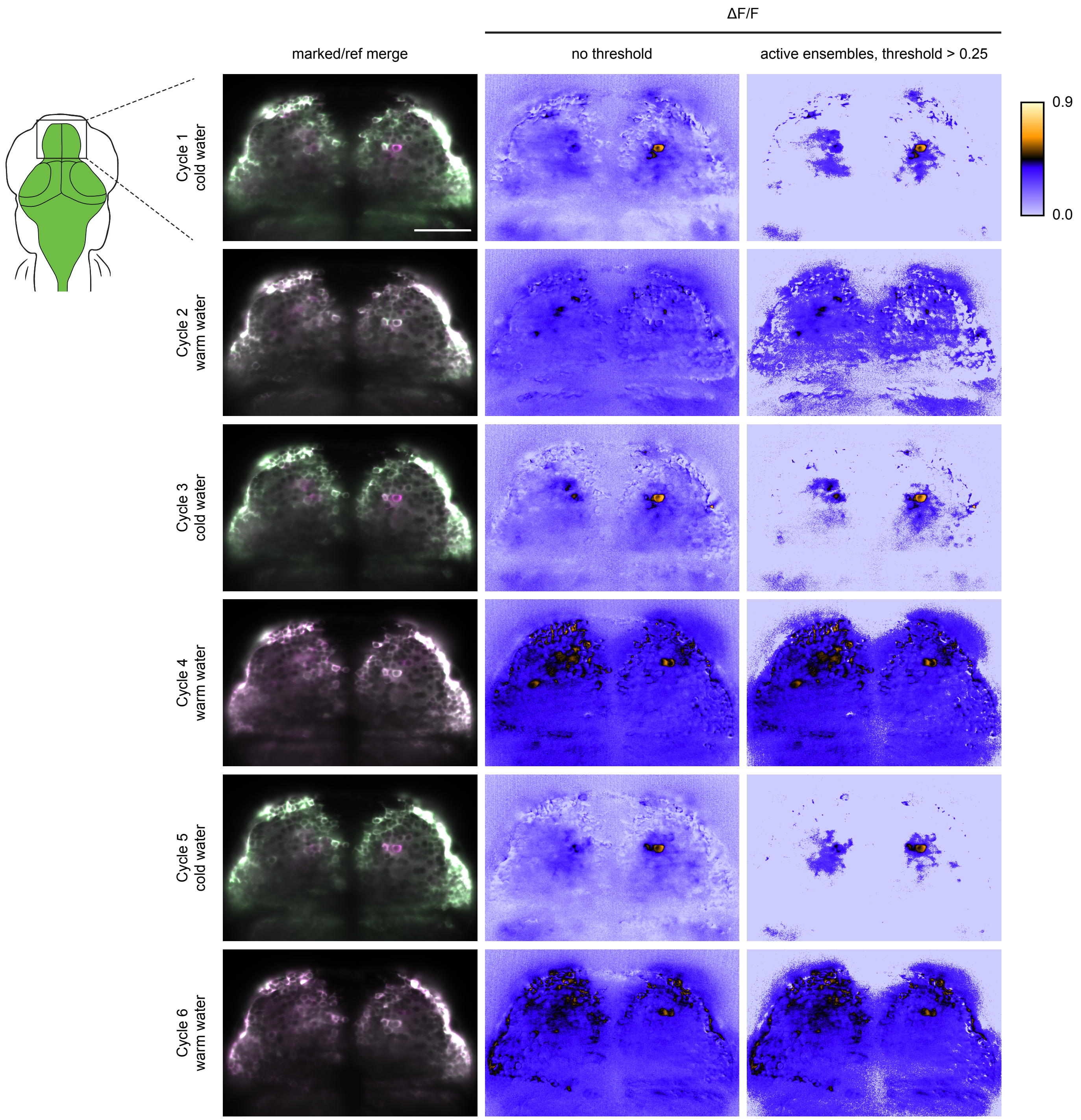


**Supplementary Figure 16: Representative Z slices of rsCaMPARI labeling from different marking cycles in the same fish.** Left image panels are merged reference (green) and erased (pseudo-colored magenta) images from the same Z position in the zebrafish pallium. Center image panels are the corresponding ΔF/F images. Right image panels are the corresponding active ensemble images with threshold above 0.25 ΔF/F. Scale bar is 50 μm.

### Supplementary Tables

| | Rel. Brightness | Initial | 490nm ↓ half-time (s) | | ↓ contrast | 405nm ↑ half-time (ms) | | ↑ contrast | Strand 8 Sequence | | Strand 9 Sequence | |
|---|---|---|---|---|---|---|---|---|---|---|---|---|
| Clone name | to rs-Eos3.1 | Contrast | + EGTA | + Ca | EGTA/Ca | + EGTA | + Ca | EGTA/Ca | N-terminal | C-terminal | N-terminal | C-terminal |
| rs-Eos3.1 | 1.00 | 1.00 | 16.93 | 14.52 | 1.17 | 229 | 220 | 1.04 | VLTGDVEMALLLE | VLTGDVEMALLLE | AHYRCDFRTTYKA | AHYRCDFRTTYKA |
| rsCaMPARI-03 | 0.57 | 1.34 | 2.11 | 6.15 | 0.34 | 500 | 504 | 0.99 | VLTGAY----SSR | TAK-TNEMALLLE |  |  |
| rsCaMPARI-17 | 0.34 | 0.35 | 24.35 | 4.28 | 5.69 | 150 | 127 | 1.18 | | | AHYRCSS---DQL | LSS-VPFRTTYKA |
| rsCaMPARI-24 | 0.24 | 0.39 | 11.79 | 2.65 | 4.45 | 139 | 130 | 1.07 | | | AHYRCDIV--DQL | LSSYTDFRTTYKA |
| rsCaMPARI-25 | 0.41 | 0.50 | 16.56 | 2.65 | 6.25 | 146 | 129 | 1.14 | | | AHYRCDFQS-DQL | LSSRTDFRTTYKA |
| rsCaMPARI-26 | 0.35 | 0.37 | 11.59 | 2.45 | 4.74 | 133 | 113 | 1.17 | | | AHYRCDGQ--DQL | LSSYTDFRTTYKA |
| rsCaMPARI-27 | 0.30 | 0.32 | 23.85 | 3.38 | 7.05 | 136 | 114 | 1.19 | | | AHYRCDKF--DQL | LSS-VPFRTTYKA |
| rsCaMPARI-29 | 0.54 | 1.16 | 1.34 | 8.10 | 0.17 | 187 | 154 | 1.22 | | | AHYRCDFYL-DQL | LSS-GTFRTTYKA |
| rsCaMPARI-30 | 0.36 | 0.35 | 5.45 | 2.76 | 1.97 | 116 | 91 | 1.27 | | | AHYRCDLG--DQL | LSS-NQFRTTYKA |
| rsCaMPARI-33 | 0.30 | 0.38 | 10.77 | 2.35 | 4.58 | 126 | 106 | 1.18 | | | AHYRCDPV--DQL | LSSSIDFRTTYKA |
| rsCaMPARI-44 | 0.46 | 0.76 | 16.19 | 4.02 | 4.02 | 131 | 132 | 0.99 | | | AHYRCDRR--SSR | TAK-AWFRTTYKA |
| rsCaMPARI-45 | 0.41 | 0.64 | 13.31 | 1.46 | 9.11 | 193 | 144 | 1.34 |  |  | AHYRDL----SSR | TAK-ALFRTTYKA |
| rsCaMPARI-46 | 0.46 | 0.64 | 13.86 | 1.22 | 11.36 | 212 | 236 | 0.90 |  |  | AHYRAL----SSR | TAK-CLFRTTYKA |
| rsCaMPARI-53 | 0.49 | 0.67 | 15.68 | 2.39 | 6.56 | 134 | 114 | 1.18 |  |  | AHYRAT----SSR | TAK-CIFRTTYKA |
| rsCaMPARI-54 | 0.62 | 1.13 | 3.05 | 16.09 | 0.19 | 168 | 169 | 0.99 | | | AHYRCLK---SSR | TAKGMDFRTTYKA |
| rsCaMPARI-61 | 0.40 | 0.62 | 17.61 | 2.29 | 7.68 | 130 | 118 | 1.11 |  |  | AHYRRE----SSR | TAK-VTFRTTYKA |
| rsCaMPARI-65 | 0.62 | 0.52 | 16.85 | 1.34 | 12.57 | 245 | 287 | 0.85 | | | AHYRCLK---SSR | TAKASDFRTTYKA |
| rsCaMPARI-67 | 0.54 | 0.66 | 16.82 | 1.63 | 10.32 | 158 | 210 | 0.75 | | | AHYRCKD---SSR | TAK---MRTTYKA |
| rsCaMPARI-78 | 0.48 | 0.75 | 12.98 | 3.28 | 3.96 | 123 | 134 | 0.92 |  |  | AHYRRG----SSR | TAK-AFFRTTYKA |
| rsCaMPARI-79 | 0.50 | 0.89 | 11.75 | 3.33 | 3.53 | 131 | 112 | 1.17 | | | AHYRCDDV--SSR | TAK-AHFRTTYKA |

**Supplementary Table 1: *In vitro* characterization of variants selected from library screening.** Nineteen selected variants from calcium-binding domain insertion screen are shown. Relative Brightness is green fluorescence normalized to mCherry red fluorescence and the template rs-Eos3.1. Initial Contrast is initial green fluorescence in $Ca^{2+}$ (0.5 mM) divided by the initial green fluorescence in EGTA (1 mM). Tabulated half-times are extrapolated from single exponential fits to the photoswitching time-course when purified protein is illuminated by 490 nm (170mW/$cm^2$) or 405 nm (200 mW/$cm^2$) light.
